# *Ex vivo* glioblastoma migration phenotypes define clinical recurrence and tumor heterogeneity

**DOI:** 10.64898/2026.08.31.748107

**Authors:** Jay Hou, Mariah McMahon, Andrew W. Ni, Zhaoning Wang, Brett Taylor, Timothy Loe, Yang Xie, Lei Chang, Andrew Gardeck, Ziya Gokaslan, Shira I. Dunsiger, Bing Ren, Clark C. Chen, David J. Odde

**Affiliations:** Department of Neurosurgery, Brown University, Providence, RI, USA; Department of Biomedical Engineering, University of Minnesota, Minneapolis, MN, USA; Warren Alpert Medical School of Brown University, Providence, RI, USA; Department of Cellular and Molecular Medicine, University of California San Diego School of Medicine, La Jolla, CA, USA; Department of Genetics and Development, Columbia University Irving Medical Center, New York, NY, USA; Division of Regenerative Medicine, Department of Medicine, University of California San Diego School of Medicine, La Jolla, CA, USA; Medical Scientist Training Program, University of California San Diego School of Medicine, La Jolla, CA, USA; New York Genome Center, New York, NY, USA; University of Minnesota Medical School, Minneapolis, MN, USA; Department of Biostatistics, Brown University, Providence, RI, USA; Department of Systems Biology, Columbia University Irving Medical Center, New York, NY, USA; Department of Biochemistry and Molecular Biophysics, Columbia University Irving Medical Center, New York, NY, USA

**Keywords:** Glioblastoma cell migration, tumor microenvironment, mechanosensitivity, gene signature, intertumoral and intratumoral heterogeneity

## Abstract

Glioblastoma’s pronounced migratory capacity underlies its diffuse invasion, presenting a formidable barrier to successful treatment. *Ex vivo* characterization of glioblastoma cells isolated from freshly resected clinical samples under physiologically relevant conditions revealed two distinct migratory phenotypes, Fast-Migrating (FM) and Slow-Migrating (SM). These phenotypes reflect distinct mechanosensitivity profiles and are associated with pharmacological responses that support the motor–clutch model of cell migration. Analysis of genes associated with these phenotypes revealed a transcriptomic signature that closely associated with *in vitro* cell migration, histological invasion in patient specimens, and clinical survival. Single-nucleus RNA sequencing revealed that FM and SM cells coexist within a single glioblastoma, with FM cells enriched at the periphery and SM cells localized to the tumor core. Collectively, our study demonstrates the utility of *ex vivo* glioblastoma characterization, allowing decoding of tumor heterogeneity and clinical prognostication as well as providing a framework for deconvoluting the complex cancer phenotype.

## Main

Glioblastoma is the most common primary malignant brain tumor in adults and remains nearly universally fatal within two years of diagnosis [1]. A key contributor to this poor prognosis involves the aggressive, infiltrative nature of the disease [2], characterized by invasion centimeters beyond the radiographically detectable tumor margins [3]. This diffuse pattern of invasion forms the basis for the current standard of care, in which radiation is delivered to a two-centimeter margin beyond the radiographically defined lesion. However, even with effective local control of the primary tumor, recurrence commonly arises outside the radiation field [4]. An understanding of the mechanisms that drive this infiltrative capacity should facilitate glioblastoma therapeutic development.

Glioblastoma exhibits significant inter-and intra-tumoral heterogeneity, which further contributes to its therapeutic resistance. Based on transcriptomic profiling, glioblastoma has been classified into distinct molecular subtypes [5, 6]. Depending on the classification scheme, these subtypes vary, reflecting underlying biological programs related to cell metabolism [7], lineage [8], and injury response [9]. While some of these biological programs are loosely associated with migratory phenotypes [10, 11], they do not fully capture the complexity of glioblastoma migration. Compounding this complexity, diverse cellular subtypes co-exist within individual tumors [12]. In this context, *ex vivo* profiling of single-cell migration may provide a functional lens for decoding these cellular complexities.

Migration, the fundamental process driving glioblastoma invasion, is critically dependent on the microenvironment’s constituents, mechanical stiffness, and the intrinsic tumor cell phenotype. The motor–clutch model of cell migration posits that actin polymerization drives membrane protrusions to form transient adhesions with the extracellular matrix (ECM), while myosin II–mediated contractility propels the cell forward [13, 14]. Each step of this process is influenced by the mechanical stiffness of the ECM [10, 15]. Magnetic resonance elastography estimates indicate that the brain’s stiffness ranges from 1 to 10 kPa (Young’s modulus), making it one of the softest tissues in the body [16–18]. Unfortunately, most conventional *in vitro* glioblastoma migration assays involve the use of rigid substrates orders of magnitude stiffer than native brain tissue [19, 20]. Moreover, most published studies rely on long-term passaged cells, which drives evolutionary drift that alters cell migration phenotypes [15, 21].

Here, we profiled the migration of glioblastoma cells isolated from 35 freshly resected clinical samples using physiologically relevant ECM ligands and substrate stiffnesses (0.7–10 kPa, Young’s modulus). Our analyses revealed two distinct migratory phenotypes (Fast-and Slow-Migrating, or FM and SM) that differ in mechanosensitivity and pharmacologic response. These phenotypes are captured by an RNA signature that correlates with histological invasion in patient specimens and clinical survival. Single-nucleus RNA sequencing revealed that FM and SM cells co-exist within a single glioblastoma, dictating bulk tumor classification based on which phenotype predominates.

### Primary glioblastoma cells exhibit distinct mechanosensitive migration phenotypes that correlate with clinical outcome

To characterize glioblastoma migration under biophysical conditions that mimic the native brain parenchyma, we profiled primary glioblastoma tumors (isocitrate dehydrogenase (IDH) wild type) from 35 patients (**Supplementary Table 1**). Freshly resected tumors were dissociated, cultured to deplete non-neoplastic cells, and evaluated within three passages to minimize *in vitro* phenotypic drift. These tumor cells were then seeded on polyacrylamide (PA) gels functionalized with brain-relevant ECM ligands (laminin, collagen I, fibronectin, and Matrigel [22, 23]). The schematic for our study design is shown in **Fig. 1a**.

**Fig. 1:**
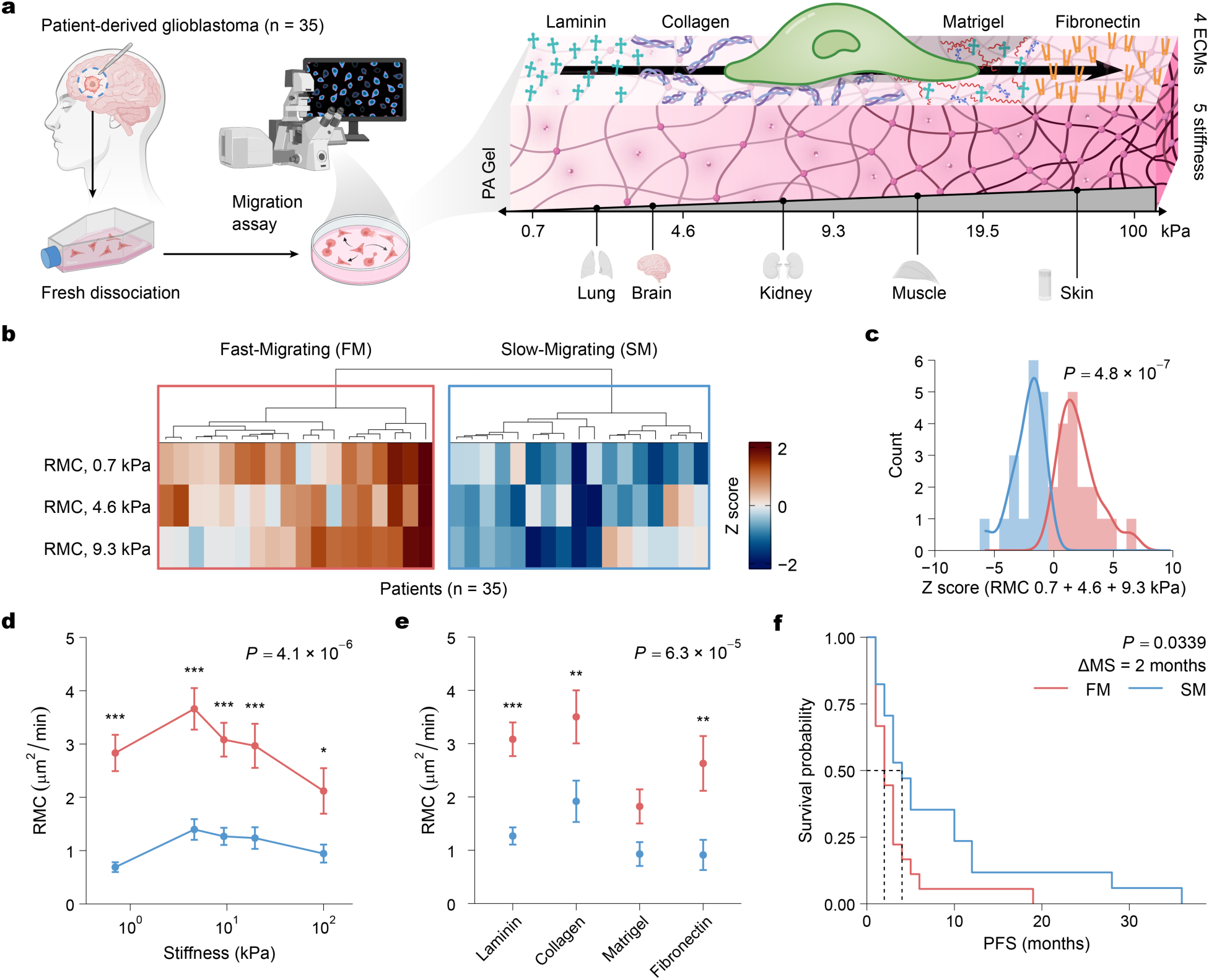
A mechanosensitive migratory phenotype in patient-derived glioblastoma cells predicts rapid tumor recurrence. **a,** Schematic of the 2D migration assay. Patient-derived glioblastoma cells (n = 35 tumors) were seeded onto polyacrylamide (PA) gels with tunable stiffness (0.7, 4.6, 9.3, 19.5, or 100 kPa), approximating the elasticity of various tissues (brain to skin) [18], and functionalized with distinct ECM ligands (laminin, collagen, Matrigel, or fibronectin). **b,** Heatmap showing hierarchical clustering (Euclidean distance, Ward’s method linkage) of patient-derived tumors (n = 35) based on random motility coefficient (RMC) quantified on substrates with stiffnesses of 0.7, 4.6, and 9.3 kPa. Clustering identifies two distinct migratory phenotypes, Fast-Migrating (FM, red) and Slow-Migrating (SM, blue). **c,** Histogram of combined RMC (0.7, 4.6, and 9.3 kPa) stratified by phenotype. The FM (red) and SM (blue) groups display significantly different distributions (*P* = 4.8×10^−7^, two-sided Wilcoxon rank-sum test). **d, e,** RMC as a function of ECM ligand (**d**) and substrate stiffness (**e**) stratified by migratory phenotype. The FM phenotype displays higher motility across all ECMs and a biphasic stiffness response (peak at 4.6 kPa) compared to the SM phenotype. Data are mean ± s.e.m. *P* values indicate the main effect of migratory phenotype (Type III ANOVA). Asterisks denote pairwise significance determined by Tukey’s post-hoc test. Significance: ns ≥ 0.05, * < 0.05, ** < 0.01, and *** < 0.001. **f,** Kaplan-Meier survival analysis of progression-free survival (PFS) reveals significantly shorter time to recurrence for the FM compared to SM group (Log-rank *P* = 0.0339, ΔMS = 2 months). ΔMS, difference in median survival months.

To identify the ECM ligand that would best support cell adherence and migration, we performed an initial screen using cells from 24 patients on ECM substrates with a stiffness of 9.3 kPa. Distinct ECM ligands supported the adherence and motility of freshly isolated cells to varying degrees (ANOVA, *P* < 0.001; **Extended Data Fig. 1a**). Although cell aspect ratio and area were comparable across substrates, laminin and collagen I supported the highest level of motility, while only laminin supported the highest level of cellular adhesion (post-hoc, *P* < 0.05, **Supplementary Fig. 1a**). Because laminin is highly enriched in the brain parenchyma [22, 23] and best supported glioblastoma adherence and migration in our screen, we selected it as the ECM ligand for subsequent experiments.

The motor–clutch model of cell migration predicts that maximal productive clutch engagement, and therefore optimal migration, occurs at an intermediate substrate stiffness [10, 15] (**Extended Data Fig. 1c**). In support of this model, pooled analysis of the cells from the full 35-patient cohort demonstrated a clear biphasic mechanosensitive response to stiffness, spanning physiological (0.7–9.3 kPa) and supra-physiological levels (19.5, 100 kPa, **Extended Data Fig. 1b, Supplementary Fig. 1b**). Cell motility (random motility coefficient, RMC) peaked at 4.6 kPa (piecewise-linear model, *P* < 0.001), which interestingly, closely reflects the stiffness of native brain tissue [16, 17], with attenuated migration at lower and higher stiffness. Similarly, cell aspect ratio (a measure of cell polarization) also peaked within this stiffness range (piecewise-linear model, *P* < 0.001). In contrast, cell area increased monotonically with stiffness (linear model, *P* < 0.01), while adherence was unaffected (**Extended Data Fig. 1b, Supplementary Fig. 1b**).

We next evaluated the migration profiles of primary cells isolated from each patient individually by applying unsupervised clustering (**Fig. 1b**). This analysis identified two distinct groups of patients with a pronounced difference in combined migration metrics (**Fig. 1c**): patients with a Fast-Migrating (FM) phenotype were characterized by high motility, polarization, and cell area regardless of substrate stiffness or ECM, while patients with a Slow-Migrating (SM) phenotype displayed the opposite traits (ANOVA, *P* < 0.001, **Fig. 1d,e, Extended Data Fig. 2)**. Notably, the FM group exhibited significant biphasic mechanosensitivity (piecewise linear model, *P* < 0.001; **Fig. 1e** and **Extended Data Figs. 2d**), characterized by a migratory peak at physiological brain stiffness (4.6 and 9.3 kPa) and enhanced motility and adherence to laminin (ANOVA, *P* < 0.01; **Fig. 1d,e** and **Supplementary Fig. 2**).

We hypothesized that progression-free survival (PFS) may serve as a proxy for migration if tumor recurrence is driven by the rate at which tumor cells escape the surgical cavity and invade the surrounding brain parenchyma. Consequently, PFS may correlate with these migratory phenotypes. Supporting this hypothesis, Kaplan-Meier analysis showed that patients with FM glioblastomas exhibited significantly shorter PFS than those with SM tumors (ΔPFS = 2 months, *P* < 0.05; **Fig. 1f**). Furthermore, this survival difference was independent of clinical covariates known to influence prognosis, as the FM/SM phenotype did not associate with age, sex, or MGMT promoter methylation (**Extended Data Fig. 3**).

### Motor–clutch mechanisms govern the biphasic response to myosin inhibition

The motor–clutch model (**Fig. 2a**) postulates that pharmacological perturbation produces predictable effects on cell migration [10]. For instance, actin inhibition is predicted to cause a monotonic decrease in motility (**Fig. 2b**) [10]. Supporting this hypothesis, actin inhibition through latrunculin A [24] uniformly decreased cell motility, aspect ratio, and area across all patients examined (n = 6; *P* < 0.01, Wilcoxon), regardless of migratory phenotype (**Fig. 2c,d** and **Supplementary Fig. 3a**).

**Fig. 2:**
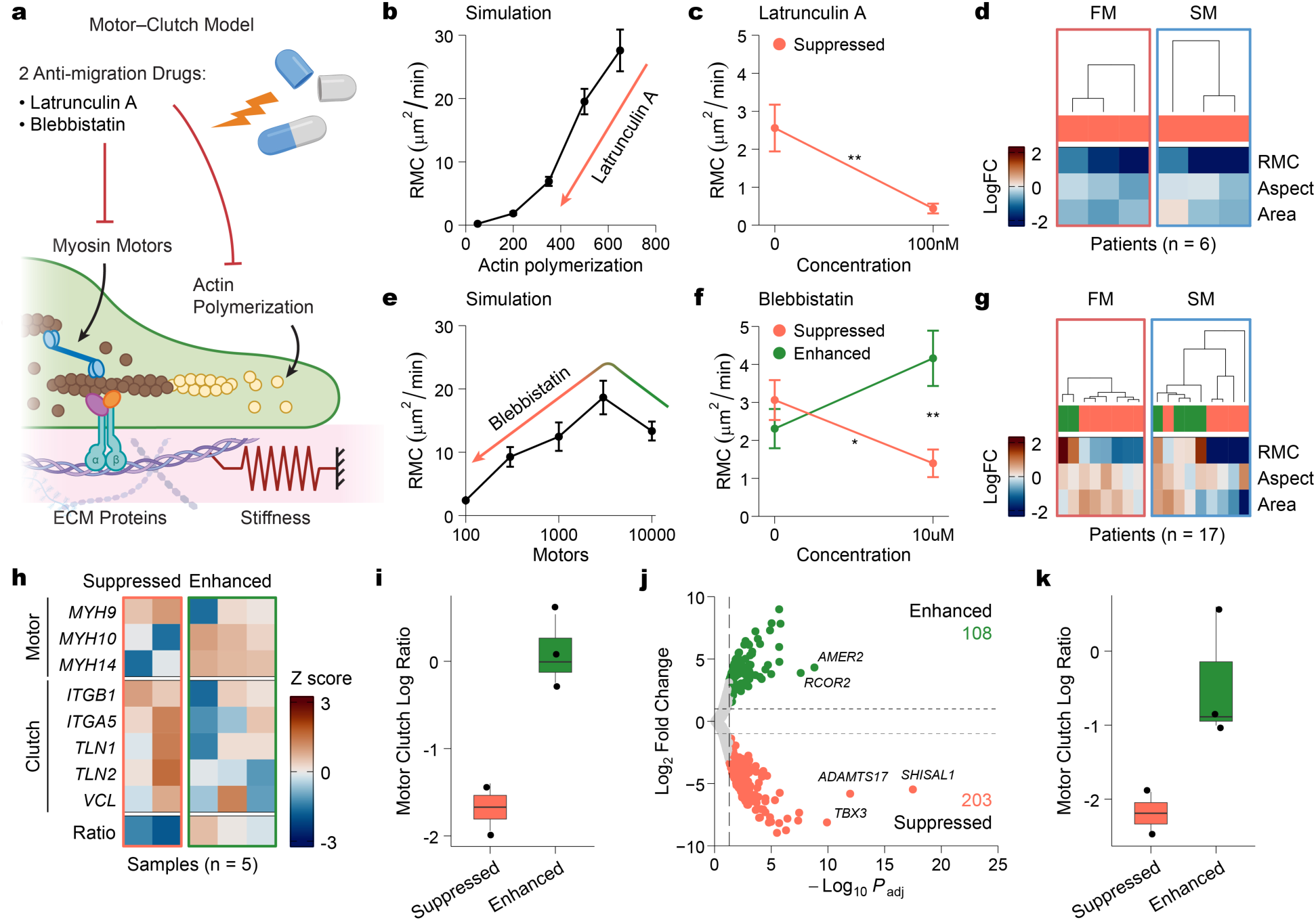
Motor–clutch dynamics predict heterogeneous migratory responses to cytoskeletal inhibition. **a,** Schematic of the motor–clutch model of cellular migration illustrating the targets of latrunculin A (actin polymerization) and blebbistatin (myosin II motors). **b,** Computational motor–clutch model simulations predict a monotonic decrease in random motility coefficient (RMC) with reduced actin polymerization. **c,** Experimental validation in patient-derived glioblastoma cells treated with latrunculin A (100 nM, n = 6 patients). **d,** Heatmap of the log2-fold change (logFC) in RMC, aspect ratio, and area for 100 nM of latrunculin A relative to control, showing consistent suppression across all patients, regardless of migratory phenotype. **e,** Simulations predict a biphasic migration response to changes in motor number (motor/clutch ratio). **f,** Experimental validation with blebbistatin treatment (10 μM, n = 17 patients) reveals some patients are enhanced (green, increased RMC) while others are suppressed (orange, decreased RMC) by blebbistatin. **g,** Heatmap (as in **d**) for blebbistatin-treated cells, showing difference between FM and SM groups in terms of enhanced and suppressed responses. **h,** Heatmap showing Z score expression of canonical motor (top) and clutch (bottom) genes in blebbistatin-suppressed and blebbistatin-enhanced samples (n = 5). **i,** Motor/clutch ratios calculated from genes in **h**, distinguishing the two groups. **j,** Volcano plot identifying differentially expressed genes (DEGs) between blebbistatin-enhanced (n = 108) and blebbistatin-suppressed (n = 203) groups (FDR < 0.05, |logFC| > 1). **k,** Motor/clutch ratios calculated from selected DEGs (**Extended Data Fig. 4d**) confirm distinct motor–clutch dynamics between the two groups. Data are mean ± s.e.m. Statistical comparisons were assessed using two-sided Wilcoxon rank-sum tests. Significance: ns ≥ 0.05, * < 0.05, ** < 0.01, and *** < 0.001.

Conversely, the motor–clutch model predicted that myosin inhibition would produce variable effects depending on the baseline motor–clutch ratio (**Fig. 2e**) [10]. Consistent with this prediction, blebbistatin [25] suppressed migration in a majority of FM cells (75%), whereas it increased migration in half of SM cells and decreased it in the other half, indicating that SM tumors span both sides of the optimal motor–clutch ratio (**Fig. 2f,g and Supplementary Fig. 3b**). These phenotype-dependent responses support the model and suggest that the enhanced migration of FM cells reflects a more optimized motor–clutch balance relative to SM cells.

To validate the motor–clutch framework, RNA sequencing (RNA-seq) was performed on five samples showing either blebbistatin-enhanced (n = 3) or blebbistatin-suppressed (n = 2) migration. Using a supervised analytical approach, we calculated motor–clutch ratios based on the expression of canonical motor (myosin) genes and clutch components (integrin, talin, and vinculin) [26, 27] (**Fig. 2h**). Supporting the motor–clutch framework, blebbistatin-enhanced tumors harbored higher baseline motor/clutch ratios than blebbistatin-suppressed tumors (**Fig. 2i**). This finding suggests that inhibition of the excessive myosin motor contributed to the increased migratory capacity of these patient cells.

Next, we tested whether an unsupervised analysis of the RNA-seq dataset would qualitatively recapitulate the findings from the supervised approach. Differential expression analysis identified 108 genes upregulated in the blebbistatin-enhanced cells and 203 genes upregulated in the blebbistatin-suppressed cells (FDR < 0.05, |log2FC| > 1; **Fig. 2j**). Pathway enrichment analysis indicated that blebbistatin-suppressed cells showed RNA expression patterns linked to negative regulation of migration, epithelial–mesenchymal transition, ECM remodeling, and integrin-mediated signaling (**Extended Data Fig. 4a,c**). In contrast, blebbistatin-enhanced tumors showed increased RNA expression related to axon guidance, differentiation, and BMP signaling (**Extended Data Fig. 4b,c**). We next identified motor and clutch genes among the differentially expressed genes and calculated motor/clutch ratios based on their expression levels. While the genes included in this unsupervised analysis differed from those used in the supervised approach, the analysis again showed that blebbistatin-enhanced tumors harbored higher baseline motor–clutch ratios than blebbistatin-suppressed tumors (**Fig. 2k and Extended Data Fig. 4d**). In aggregate, the above analysis suggests that the intrinsic motor–clutch state of primary glioblastoma cells determines their response to myosin inhibition.

### A 13-gene migration signature captures the migratory phenotypes and associates with clinical outcomes

Our results suggest that the migratory behavior of glioblastoma is, at least in part, intrinsically encoded at the transcriptional level. To identify the transcriptional programs that define the FM group, we analyzed RNA-seq data from 18 patient-derived glioblastoma lines with characterized migratory phenotypes [10] (**Fig. 3a**). Based on the combined RMC Z score across 0.7, 4.6, and 9.3 kPa substrates (**Extended Data Fig. 5a**), the patient cell lines were classified into FM and SM groups (**Fig. 3b**). Principal component analysis of the transcriptomes partitioned the tumors in alignment with their migratory classification, confirming that distinct cellular programs contribute to the FM and SM phenotypes (**Extended Data Fig. 5b,c**).

**Fig. 3:**
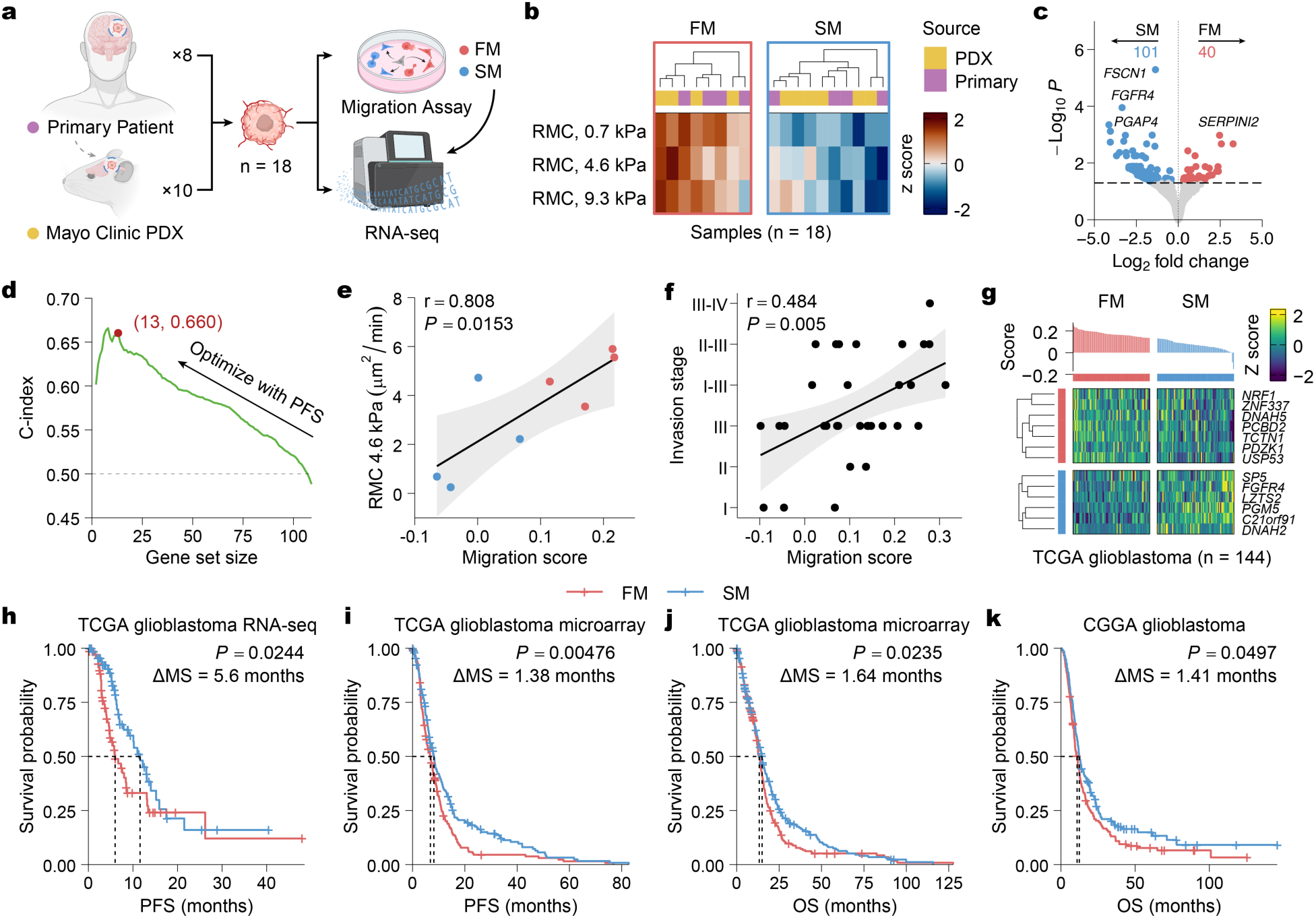
A gene signature capturing the migratory phenotypes predicts motility and clinical prognosis. **a,** Schematic of the gene signature discovery workflow combining RNA-seq and 2D migration assays. Analyses were performed on primary patient (n = 8) and Mayo clinic PDX samples (n = 10). **b,** Heatmap showing separation of samples (n = 18) into migratory phenotypes, Fast-Migrating (FM) (red) and Slow-Migrating (SM) (blue) (see **Extended Data Fig. 5a**). **c,** Volcano plot showing DEGs between FM (n = 40) and SM (n = 101) phenotypes (FDR < 0.05). **d,** Stepwise optimization of the migration gene signature using the TCGA RNA-seq cohort (n = 144, IDH1-mutant excluded). DEGs were iteratively removed to maximize Harrell’s concordance index (c-index) for PFS, yielding an optimized 13-gene signature (c-index = 0.66). **e,** Prospective validation of the migration signature to predict *in vitro* cell motility (random motility coefficient, RMC) on 4.6 kPa substrates (Pearson r = 0.808, *P* = 0.0153). **f,** Correlation of migration score with *in vivo* histological invasion stage (Pearson r = 0.484, *P* = 0.005) across a larger cohort of Mayo PDX lines (n = 32). Shaded areas in **e** and **f** indicate the 95% confidence intervals. **g,** Distribution of enrichment scores (top) and expression heatmap of the 13-gene migration signature (bottom) in the TCGA RNA-seq cohort (n = 144, IDH1-mutant excluded), stratifying patients by median score. **h–k,** Kaplan-Meier survival analysis of FM (red) and SM (blue) groups stratified by median migration score. The migration signature predicts significantly shorter PFS in the TCGA glioblastoma RNA-seq cohort (**h**, n = 144, ΔMS = 5.6 months, *P* = 0.0244) and microarray cohort (**i**, n = 525, ΔMS = 1.38 months, *P* = 0.00476), and shorter overall survival (OS) in the TCGA glioblastoma microarray cohort (**j**, n = 525, ΔMS = 1.64 months, *P* = 0.0235) and CGGA glioblastoma cohort (**k**, n = 298; IDH1-mutant excluded, ΔMS = 1.41 months, *P* = 0.0497). *P* values were determined by two-sided log-rank test. ΔMS, difference in median survival months.

Differential expression analysis between FM and SM cell lines identified 141 differentially expressed genes (FDR < 0.05; **Fig. 3c**). Pathway analysis [28] revealed that the SM cell lines harbored RNA expression associated with ECM organization [22], substrate adhesion [29], and bone morphogenic protein (BMP) signaling [30] (**Extended Data Fig. 6a**). In contrast, the FM cell lines harbored RNA expression associated with actin-rich cell projections [31], ciliogenesis and axoneme function [32, 33], and glycogen metabolism [34] (**Extended Data Fig. 6b**). These results indicate that FM cells harbor transcriptional programs that facilitate mechanosensitivity and migration, whereas SM cells are enriched for pathways involved in ECM reorganization. Using a network-based method [28] to cluster the enriched pathways provided further support for this hypothesis, revealing two transcriptional modules: FM characterized by an actin–microtubule–metabolic program, and SM defined by an ECM–adhesion–BMP program (**Extended Data Fig. 6c**).

As a first step toward a clinically actionable biomarker, we refined the differentially expressed genes into a migration gene signature by optimizing concordance with PFS using patients from The Cancer Genome Atlas (TCGA) glioblastoma RNA-seq dataset (n = 144, IDH1-mutant excluded, [35]) (**Fig. 3d**). This iterative optimization process yielded a 13-gene migration signature comprising seven genes upregulated in the FM state (NRF1, ZNF337, DNAH5, PCBD2, TCTN1, PDZK1, USP53) and six upregulated in the SM state (SP5, FGFR4, LZTS2, PGM5, C21orf91, DNAH2). Robustness analysis using bootstrap resampling demonstrated that the signature outperformed 99% of randomly sampled gene sets across multiple survival metrics (**Extended Data Fig. 7**).

Using this signature, we computed the migration score for the original patient lines using single-sample gene set enrichment scoring with singscore [36]. Despite comprising far fewer genes, the migration signature reproduced the FM and SM phenotypes, correctly classifying 100% of the original patient lines (**Extended Data Fig. 5d**). Moreover, the signature scores strongly correlated with experimentally measured motility in the 18 lines (r = 0.766, *P* < 0.001; **Extended Data Fig. 5e**). To validate the 13-gene signature as a biomarker of glioblastoma migratory capacity, we prospectively characterized migration in eight previously untested patient-derived glioblastoma lines and calculated their corresponding signature scores, all performed in a blinded manner to minimize bias. Upon unblinding of the data, we found that the 13-gene score showed a strong correlation with experimentally measured motility (r = 0.808, *P* < 0.05; **Fig. 3e**), correctly classifying FM/SM phenotype in 75% (6/8) of the glioblastoma lines studied using a median split (**Extended Data Fig. 5f**).

Given that migration is a fundamental component of tumor invasion, we next assessed whether the migration signature correlated with histological invasion in 32 previously published clinical glioblastoma specimens with matched RNA-seq data. Invasion scores were previously assigned [37], and a 13-gene signature score was calculated for each sample using the available RNA-seq data. Impressively, higher migration scores were associated with higher histological invasion stages (r = 0.484, *P* < 0.01; **Fig. 3f**). These findings suggest that the transcriptional programs captured by the 13-gene signature reflect the biological processes that drive glioblastoma invasion.

The 13-gene signature was optimized based on its concordance with PFS in the TCGA RNA-seq dataset (n = 144). Expectedly, when patients in this cohort were stratified into FM and SM groups using the median migration score (**Fig. 3g**), their survival outcomes differed significantly (ΔMS = 5.6 months, *P* < 0.05; **Fig. 3h**). Notably, migration score remained the only independent, adverse predictor of PFS in multivariate Cox regression after adjusting for confounders, including MGMT methylation, molecular subtype, age, and sex (*P* < 0.01; **Supplementary Fig. 4a**).

To further validate the clinical association of the 13-gene signature, we evaluated it using the TCGA microarray dataset (Firehose Legacy, n = 525), acknowledging the limitation that the patients in this dataset included some of the 144 patients in the RNA-seq cohort. Using the same stratification strategy, we found patients in the FM group exhibited shorter PFS relative to those in the SM group (ΔMS = 1.38 months, *P* < 0.01; **Fig. 3i**). Given that PFS is a key determinant of overall survival (OS), we also tested whether the 13-gene signature associated with OS using the TCGA microarray dataset. When the 525-patient cohort was classified into FM or SM groups using the signature, FM patients exhibited significantly shorter OS (ΔMS = 1.64 months, *P* < 0.05; **Fig. 3j**).

As a final validation effort, we turned to an entirely different dataset, the Chinese Glioma Genome Atlas (CGGA; n = 298, IDH1-mutant excluded, [38]). In this independent cohort, FM patients again exhibited a shorter OS relative to the SM patients when stratified by median migration score (ΔMS = 1.41 months, *P* < 0.05; **Fig. 3k**). Moreover, migration score, in addition to tumor type and age, remained an independent adverse predictor of OS in multivariate Cox regression (*P* < 0.05; **Supplementary Fig. 4b**). These findings suggest that the 13-gene signature may bear clinical prognostic utility by capturing transcriptional programs associated with glioblastoma migration.

### The migration signature is orthogonal to established subtype signatures

Transcriptomic analyses have classified glioblastoma into multiple molecular subtypes [5, 6], including those reflecting cell lineage [8], developmental and injury-response programs [9], and metabolic states [7] (**Fig. 4a**). However, these frameworks do not fully capture the biology of cell migration. We therefore hypothesized that migration-associated transcriptional programs may overlap with, yet remain independent of, established subtype classifications. Supporting this hypothesis, the 13-gene migration signature showed no overlap with any previously published subtype classifiers (Jaccard index = 0; **Fig. 4a**), and patient stratification by migration score did not align with established subtypes in the TCGA glioblastoma RNA-seq cohort (n = 144, IDH1-mutant excluded) (**Extended Data Fig. 8a–d**).

**Fig. 4:**
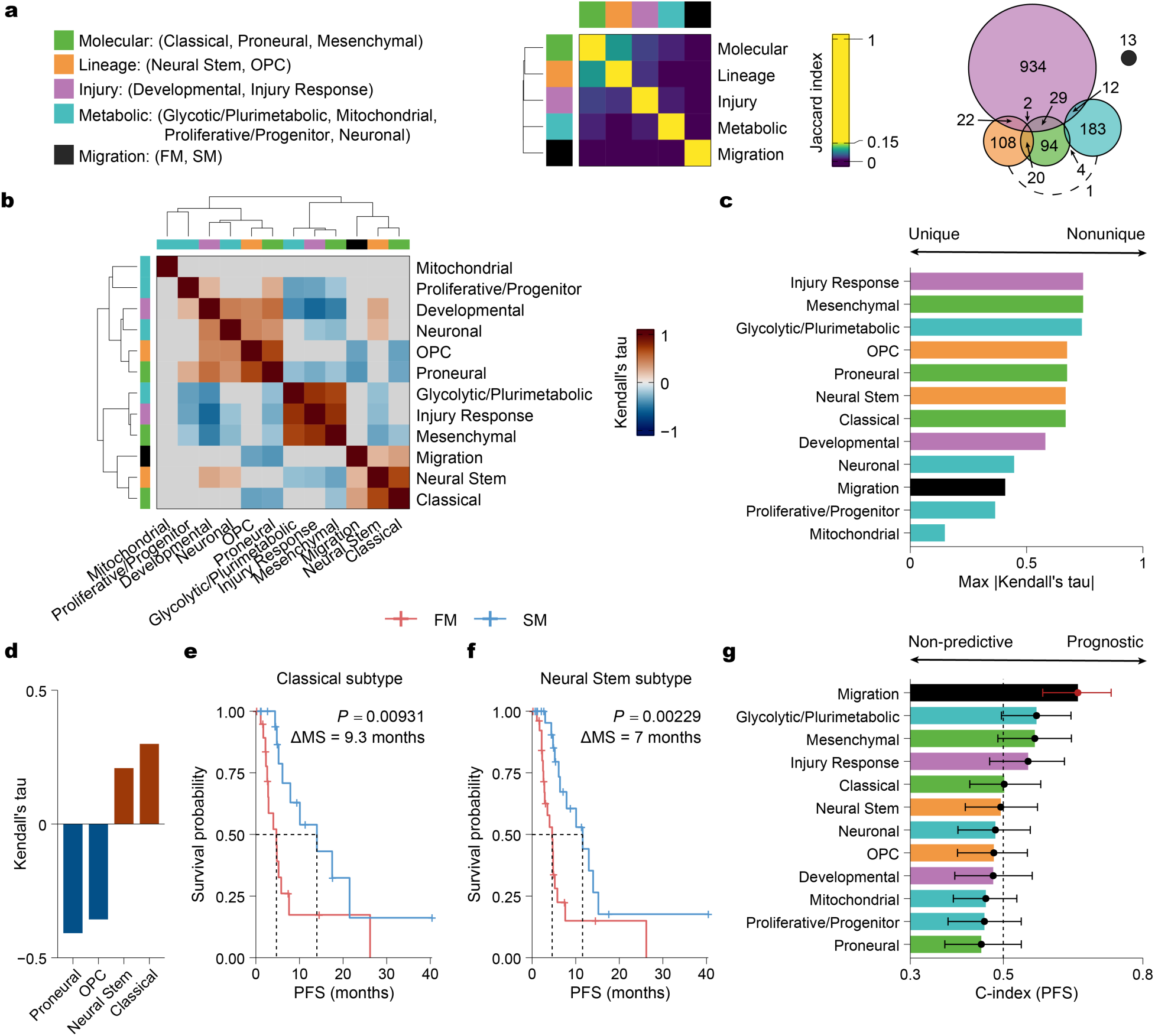
The migration signature captures a prognostic axis orthogonal to established glioblastoma subtypes. **a,** Heatmap of Jaccard indices (left) and Venn diagram (right) quantifying gene set overlap between the migration signature and established glioblastoma signatures, including molecular subtypes (green) [5, 6], cell-lineage (orange) [8], developmental/injury response programs (purple) [9], and metabolic states (blue) [7]. The migration signature (black) shares no genes with these established signatures. **b,** Pairwise concordance of patient progression free survival (PFS) rankings for each signature in the TCGA glioblastoma RNA-seq cohort (n = 144, IDH1-mutant excluded), assessed by Kendall’s tau. Gray cells indicate non-significant associations (Bonferroni-corrected *P* > 0.05). **c,** Signature redundancy analysis comparing maximum absolute Kendall’s tau for each signature against all others. The migration signature exhibits low maximal tau, indicating it captures variation independent of other subtypes. **d,** Correlation (Kendall’s tau) of the migration signature with specific subtypes, highlighting significant positive (e.g., Classical) and negative (e.g., Proneural) associations (*P* < 0.05). **e, f,** Kaplan–Meier survival analysis within positively correlated subtypes in **d** shows significantly shorter PFS in the FM versus SM group (quartile split) for a subset of patients with Classical (**e**, n = 78/144, ΔMS = 9.3 months, *P* = 0.00931) and Neural Stem tumor subtypes (**f**, n=108/144, ΔMS = 7 months, *P* = 0.00229). **g**, Prognostic accuracy of each subtype for PFS in the TCGA glioblastoma RNA-seq cohort, assessed by c-index. The migration signature (black) demonstrates superior prognostic ability compared to established subtypes. Error bars represent 95% confidence intervals from bootstrap resampling. Red points denote subtypes where this interval excludes the null value of 0.5.

To quantify the relationship between established subtype signatures and our 13-gene migration signature, we assessed concordance in patient rankings within the TCGA RNA-seq cohort using Kendall’s tau [39]. While established subtypes displayed strong pairwise correlations, the migration signature did not, exhibiting the lowest maximum tau values (**Fig. 4b**,**c**). Among these comparisons, the migration signature was modestly negatively correlated with the proneural and oligodendrocyte precursor (OPC) subtypes and positively correlated with the classical and neural stem signatures (**Fig. 4d, Extended Data Fig. 8a–d**).

These analyses demonstrate that the migration signature is largely orthogonal to established molecular subtypes, capturing transcriptional variation not entirely accounted for by lineage, developmental, or metabolic programs. Notably, even within positively correlated subtypes, the migration score was still able to stratify patients into FM and SM groups with significantly different PFS, for both classical (ΔMS = 9.3 months, *P* = 0.00931) and neural stem subtypes (ΔMS = 7 months, *P* = 0.00229) (**Fig. 4e,f**). Moreover, the migration signature remained the only significant prognosticator of PFS among all evaluated subtype signatures (**Fig. 4g; Extended Data Fig. 8e**).

### Intra-tumoral migratory heterogeneity in glioblastoma

Given the pronounced intratumoral heterogeneity of glioblastomas [5], we next examined whether the migration gene score, reflecting tumor cell migratory capacity, mapped to distinct spatial regions within the tumor. To this end, we performed single-nucleus RNA sequencing (snRNA-seq) on clinical glioblastoma samples collected from the tumor core (non-contrast enhancing), tumor edge (enhancing rim), and adjacent normal cortex (non-enhancing tissue beyond the tumor margin) of four glioblastoma patients [40] (**Fig. 5a**). After identification of malignant cells using copy number variation (CNV) inference (**Extended Data Fig. 9a–e**), we calculated single-cell migration scores using our 13-gene signature (**Fig. 5b**). We classified cells as either FM or SM based on migration signature Z scores (**Fig. 5c**). Strikingly, FM cells were strongly enriched at the tumor edge and within adjacent normal brain tissue, whereas SM cells were localized to the tumor core (**Fig. 5d and Extended Data Fig. 9f,g**). This established a distinct outward gradient of increasing migratory capacity from the inner core to the invasive margin.

**Fig. 5:**
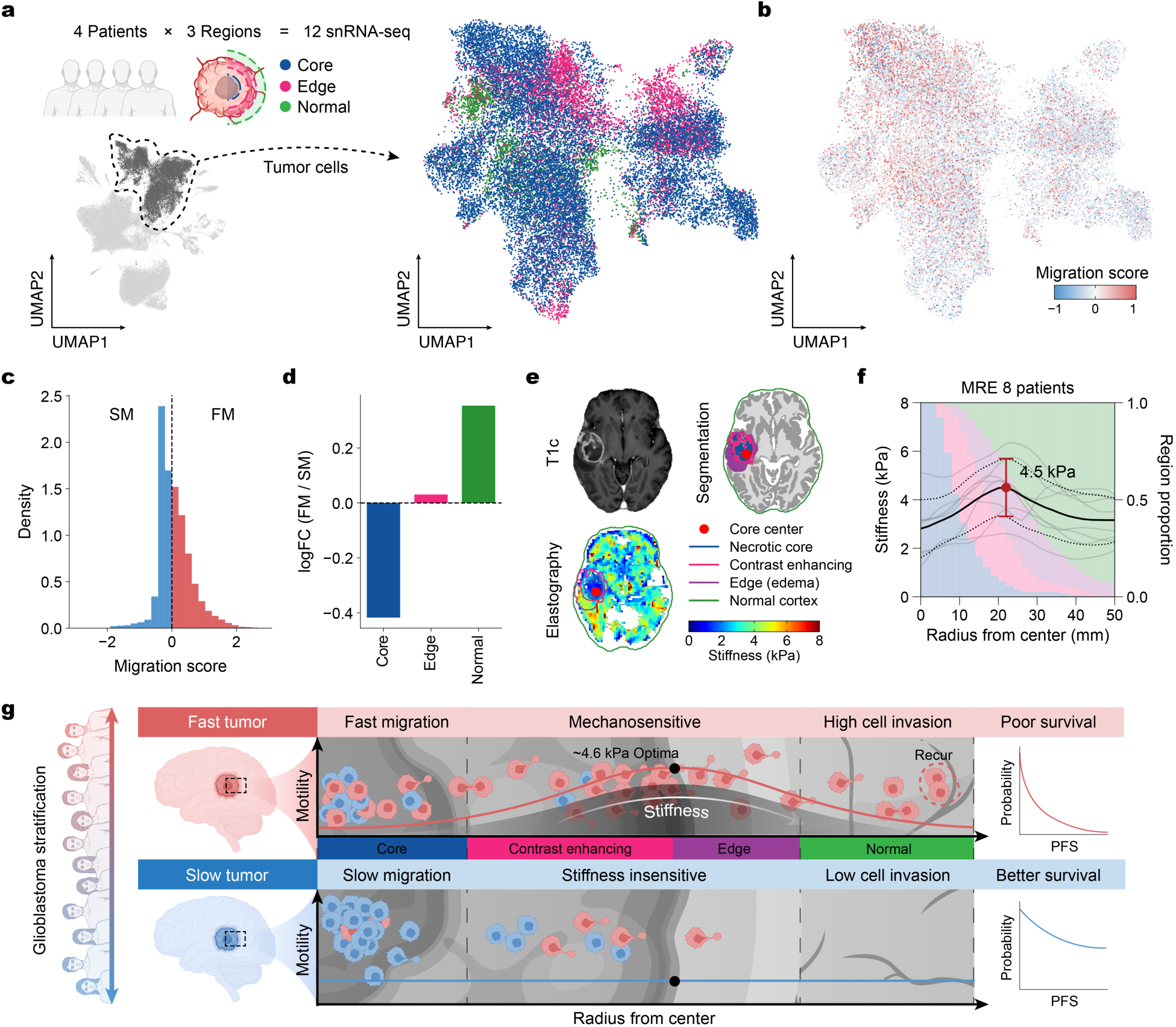
Fast, mechanosensitive glioblastoma cells accumulate at the stiff tumor edge to drive invasion. **a,** Schematic of the snRNA-seq sampling strategy (left) analyzing 12 samples from 4 glioblastoma patients across 3 spatial regions: tumor core (blue), edge (pink), and radiographically normal brain (green). Tumor cells were identified (left UMAP, dashed line outlines tumor cluster) and analyzed (right UMAP). **b,** UMAP projection of per-cell migration scores derived from the migration signature. Warmer colors indicate higher migration scores (FM-like), and cooler colors indicate lower scores (SM-like). **c,** Distribution of migration scores across all tumor cells, showing the threshold (dashed line at 0) used to classify cells as Fast-Migrating (FM) or Slow-Migrating (<0). **d,** Log2-fold change (logFC) of the FM/SM cell ratio across regions demonstrates progressive enrichment of FM cells in the edge and normal regions compared to the core. **e,** Representative patient imaging showing a co-registered T1c scan (top left), tumor segmentation defining the necrotic core, contrast-enhancing rim, edema (edge), and normal cortex (top right), and the corresponding magnetic resonance elastography (MRE) stiffness map (bottom left). The red dot indicates the tumor center. **f,** MRE glioblastoma stiffness profile (n = 8 patients). Background colors represent the proportion of each tissue region (gray, necrotic core; pink, contrast enhancing; blue, edema; green, normal cortex) at a given radial distance. Peak stiffness (Young’s modulus 4.5 kPa) coincides with the transition from tumor edge to normal tissue and is strikingly close to the biphasic peak (4.6 kPa) of fast, mechanosensitive cells. **g,** Proposed framework for glioblastoma stratification and a model of mechanosensitive, glioblastoma invasion. Tumors enriched for the fast signature (top) have more fast, mechanosensitive cells which preferentially migrate along the stiffness gradient to an optimum of ∼4.6 kPa at the tumor edge, leading to high cell invasion, recurrence, and poor outcomes. Conversely, tumors enriched for the slow signature (bottom) have more slow, stiffness-insensitive cells with limited motility, resulting in less cell invasion and better survival outcomes.

We validated this finding in an independent, similarly spatially annotated single-cell RNA-seq (scRNA-seq) dataset from a glioblastoma patient undergoing temporal lobectomy with intraoperative 5-aminolevulinic acid (5-ALA) fluorescence guidance [41]. After identifying malignant cells via CNV profiling, we again classified them into FM and SM subpopulations based on their migration scores (**Extended Data Fig. 10a–c**). Consistent with our initial analysis, regional analysis in this independent cohort again showed FM cells enriched at the tumor periphery and in normal cortex, whereas SM cells were concentrated in the necrotic core and high-fluorescence tumor regions (**Extended Data Fig. 10d–f**). Together, these data suggest that cells with high migratory capacity are preferentially localized to the cerebrum adjacent to the tumor edge.

To relate this spatial organization to the mechanical microenvironment and the mechanosensitivity we observed in the FM phenotype, we analyzed co-registered MRI and magnetic resonance elastography (MRE) scans from eight glioblastoma patients [17] (n = 8; **Fig. 5e**). Integrating tumor segmentation with spatial stiffness measurements revealed a remarkably consistent radial stiffness gradient across regions (**Supplementary Fig. 5a–i**). The necrotic cores were relatively soft (∼3 kPa, equivalent Young’s modulus), whereas stiffness increased across a peritumoral transition zone, peaking at approximately 4.5 kPa at the tumor edge (**Fig. 5f, Supplementary Fig. 5j,k**). Strikingly, this peak in stiffness aligns almost exactly with the optimal stiffness (4.6 kPa) required for maximal glioblastoma cell motility we previously discovered in FM tumors (**Fig. 1d**).

Finally, the co-existence of FM and SM cells prompted us to revisit our initial data with the hypothesis that the classification of tumors as FM or SM (**Fig. 1b**) reflects the relative abundance of these phenotypes within each tumor. Using the median RMC to classify each cell within the tumor as FM or SM, we found that both populations co-exist within the same tumor (**Supplementary Fig. 6a,b**). Supporting our hypothesis, glioblastomas classified as FM harbored a higher proportion of FM cells, whereas SM glioblastomas were enriched for SM cells (**Supplementary Fig. 6b,c**). These findings indicate that the bulk migratory phenotype of a glioblastoma is largely determined by the relative composition of FM and SM populations.

Together, these data support a mechanosensitive invasion model for glioblastoma in which FM cells accumulate at the stiff tumor edge, enabling their highly migratory phenotype to drive parenchymal infiltration and seed clinical recurrence (**Fig. 5g**).

## Discussion

Glioblastoma’s aggressive migratory capacity drives diffuse infiltration, compromising therapeutic efficacy and patient survival [2]. To characterize the molecular underpinnings of this infiltration, we examined the migratory phenotype of freshly dissected glioblastoma cells from patient-derived specimens using physiologically relevant ECM substrates and stiffness. Our study revealed two distinct migratory modes: Fast-Migrating (FM) and Slow-Migrating (SM). FM cells display biphasic, mechanosensitive migration that peaks at native brain stiffness. In contrast, SM cells are insensitive to substrate stiffness, maintaining a uniformly low migratory speed across mechanical conditions. Latrunculin A uniformly suppresses the migratory speed of all cells, while the effect of blebbistatin on migration varies depending on their baseline motor–clutch balance. The differences between these phenotypes are captured by a transcriptomic signature that correlates robustly with *in vitro* migration, histopathological invasion in patient specimens, and clinical survival. Single-nucleus RNA sequencing further revealed that FM and SM cells co-exist within the same tumor, with overall tumor classification reflecting their relative abundance. Spatially, FM cells were enriched at the invasive tumor edge, where tissue stiffness is optimal to support FM migration, whereas SM cells were more frequently localized within the tumor core.

Our finding that myosin II inhibition with blebbistatin can paradoxically enhance the migratory speed of glioblastoma cells suggests that non-discriminatory targeting of the cellular motor apparatus may inadvertently promote diffuse infiltration and potentiate disease progression in a subset of patients. The identified 13-gene transcriptomic signature provides a potential framework for rational patient stratification, ensuring that motor–clutch-targeted therapies are deployed only in individuals with glioblastoma harboring susceptible phenotypes. Another viable translational strategy involves targeting F-actin assembly rather than contractility, since latrunculin A uniformly suppresses the migratory speed of all glioblastoma cells. For example, the available literature increasingly highlights the F-actin-bundling protein fascin, which coordinates the formation of sensory filopodia at the invasive leading edge, as a promising anti-migratory target [42]. Notably, the antidepressant imipramine has recently been shown to inhibit fascin [43], raising the possibility of its repurposing as a therapeutic agent in glioblastoma [44, 45].

The 13-gene signature that stratifies glioblastoma into FM and SM phenotypes exhibits limited overlap with existing transcriptomic schemes, demonstrating that this migratory axis is distinct from underlying subtype programs. Supporting this hypothesis, the 13-gene signature robustly captures histopathological invasiveness and patient survival beyond the predictive capacity of available subtype signatures [5–9]. If prospectively validated, this transcriptomic tool could directly guide surgical decision-making. Given the tendency for diffuse invasiveness in FM-dominated tumors, aggressive cytoreductive surgery is unlikely to extend patient survival meaningfully and may instead expose frail individuals to surgical or perioperative risks. Consequently, screening vulnerable or elderly populations with this signature can help surgeons balance therapeutic utility against potential morbidity. Beyond surgical stratification, this orthogonal classification provides a valuable platform for stratified clinical trial enrollment, longitudinal monitoring of phenotypic evolution, and the rational design of targeted, anti-invasive combination therapies.

The intratumoral heterogeneity in migratory behavior in glioblastoma supports a model in which intrinsic migratory programs interact with the mechanical microenvironment to shape invasion. These intrinsic programs are governed by motor and clutch proteins, whose expression generally does not change in response to differences in substrate stiffness [46]. Consistent with the role of intrinsic programs, FM cells are observed even in tumor regions where ECM stiffness is less optimal for migration (**Fig. 5a,b**). At the same time, the preferential enrichment of FM cells at the tumor periphery, together with *in vitro* findings that glioblastoma cells migrate durotaxically along stiffness gradients [46] suggest that the enrichment of FM cells at the tumor periphery may reflect the convergence of an intrinsic migratory state with a mechanically permissive microenvironment (**Fig. 5g**). The relative contributions of cell-intrinsic programs and extrinsic mechanical signals to the migration phenotype remain an open question.

The classification of migratory phenotypes into FM and SM, however, likely oversimplifies a continuous spectrum of cell behaviors. As sample sizes increase and more diverse microenvironmental contexts are examined, additional thresholds or migratory states may emerge. Moreover, identifying RNA signatures across substrates with varying stiffness, substrates, or within alternative migratory environments, such as organoid systems [47], could provide insights into other aspects of glioblastoma migration. For instance, Wong et al. [20], using microfluidic systems with relatively stiff substrates (∼1MPa), likely captured migratory programs engaged under mechanical conditions that differ substantially from those found in the brain, contributing to an RNA signature distinct from ours (**Supplementary Fig. 7**). Together, these studies provide a proof of principle for linking migration phenotypes with transcriptional states, while underscoring the need for future work that integrates diverse microenvironmental contexts.

In sum, *ex vivo* characterization of glioblastoma cells from freshly resected specimens revealed migratory phenotypes that are associated with distinct cellular programs, pharmacologic responses, and clinical outcomes. This mechanism has broad relevance across other solid tumor types where mechanical stiffness drives malignancy. Finally, our study provides a blueprint for translating single-cell biophysical behaviors into personalized oncology strategies.

## Methods

### Patient-derived glioblastoma cell culture

Primary tumor samples were collected by C.C.C. with informed consent, from 35 patients undergoing surgical resection for glioblastoma at the University of Minnesota Medical Center (2017–2019; IRB approval: 9702M12335, STUDY00001136, IRB 120345X). Samples were transported on ice and processed within 16 h to preserve viability. Tumors were mechanically dissociated, washed in PBS (Gibco 10010-023), and treated with ACK lysis buffer (Gibco LSA1049201) to remove red blood cells. Cells were centrifuged and resuspended in neural stem cell media consisting of DMEM/F-12 (Gibco 11320033) supplemented with 1× B-27 (Gibco 12587-010) and 1× penicillin–streptomycin (Corning 45000-650). The suspension was filtered through a 40 µm strainer and plated on Matrigel-coated flasks (Corning 354263) with EGF (Peprotech AF-100-15) and FGF (Peprotech AF-100-18B). Cultures were maintained with media changes every 48 h until debris and immune cells were depleted (3–10 days) prior to imaging. Only patient lines with sufficient high-quality trajectories (≥30 cells per condition) were included in the study.

### Polyacrylamide hydrogel synthesis

Polyacrylamide (PA) gels spanning five stiffness levels (nominal Young’s moduli of 0.7, 4.6, 9.3, 19.5, and 100 kPa) were prepared as previously described [15]. Briefly, precursor solutions of 40% acrylamide (Fisher BP1402), 2% bis-acrylamide (Fisher BP1404), 1 M HEPES (Sigma H6147), and deionized water were mixed to achieve the desired stiffness. After degassing, solutions were supplemented with 1% (v/v) ammonium persulfate (Bio-Rad 161-0700) and TEMED (Fisher BP150), cast onto silanized glass-bottom dishes (MatTek P35G-0-20-C), and covered with hydrophobic coverslips (Fisher 12-545-80) to form ∼100 µm thick gels. Following polymerization, the PA gels were activated with sulfo-SANPAH (Thermo 22589) under UV illumination and functionalized overnight at 4 °C with the desired ECM protein: laminin (Corning 354232), Matrigel (Corning 354230), collagen I (Corning 354236), or fibronectin (Sigma-Aldrich F1141).

### Migration assay

Glioblastoma cells were plated on PA gels and allowed to adhere overnight. Time-lapse microscopy was performed as previously described [15]. Multi-position phase-contrast images were acquired every 15 min for 12 h using a Nikon Eclipse TE200 microscope equipped with a Plan Fluor 10×/0.30 NA objective and automated XY stage (NIS-Elements AR v4.5). Image processing and cell tracking were conducted using custom MATLAB scripts [10]. Raw image stacks were flat-field corrected and background-subtracted. Binary masks were generated via adaptive thresholding, followed by morphological opening/closing, hole filling, and watershed segmentation. Object features, including centroids, area, and major/minor axes, were extracted for each frame and linked across frames using the Hungarian algorithm. From the resulting trajectories, we computed the random motility coefficient (RMC), aspect ratio (major/minor axis), area, and adherence (fraction of cells remaining attached throughout the 12 h imaging period). RMC was derived from mean squared displacement using the 2D diffusion model as previously described [15]. A subset of the cell migration data has been previously published [48].

### Pharmacological inhibition

To investigate the specific roles of cytoskeletal dynamics in cell motility, migration inhibitors were utilized. For actin disruption, cells were treated with latrunculin A (Sigma-Aldrich 428021-100UG; 100 nM). For myosin II inhibition, cells were treated with blebbistatin (Sigma-Aldrich B0560-1MG; 10 µM). Doses were selected based on their lack of measurable toxic effects on cells [24, 49]. Drugs were dissolved in dimethyl sulfoxide (DMSO) to create stock solutions and diluted in culture media immediately prior to use. Control conditions received an equivalent volume of DMSO vehicle (final concentration <0.1% v/v). Treatments were applied 2 h prior to the start of microscope imaging and maintained throughout the acquisition period. Trajectories were analyzed as described previously, and comparisons were made between drug-and vehicle-treated cells.

### Computational motor–clutch modeling

The governing equations and algorithms of the Cell Migration Simulator (CMS) have been described previously [10, 15]. Briefly, the CMS represents a cell as multiple protrusive modules that are nucleated stochastically at a rate determined by the maximum nucleation rate k_mod and the total actin pool a_total. Protrusions elongate at a polymerization rate determined by v_poly and a_total. Protrusions are stochastically capped at rate k_cap, terminating elongation, and are removed if their length falls below a minimum threshold l_min.

Cell movement is determined by force balance between protrusive module forces and the cell body force across modules. Forces arise from a motor–clutch framework, in which individual clutch forces depend on the clutch spring constant k_c and clutch displacement. The total force contributions from the cell body and protrusive modules are given by the sum of individual clutch forces across n_cell cell-body clutches and n_c protrusion clutches. Clutches bind to F-actin with rate k_on and unbind with a rate that depends on a baseline unbinding rate k_off, the force on the clutch, and the motor force F_m, where F_m is given by the product of motor number n_m and single-motor force F_m.

Parameter values were chosen based on previous simulations (**Supplementary Table 2**)[10]. Simulations were performed using a Monte Carlo framework based on the Gillespie stochastic simulation algorithm, in which events were selected probabilistically according to cumulative rates and time steps were determined from the total rate and a uniform random number. The C++ implementation of the CMS [10] was used for all simulations and run on the Mesabi cluster at the Minnesota Supercomputing Institute (MSI). To simulate myosin inhibition (blebbistatin), the motor–clutch ratio was modified by reducing the number of motors n_m while keeping the number of clutches n_c constant. To simulate actin disruption (latrunculin A), the actin polymerization rate v_poly was systematically decreased.

### Migration data analysis

#### Preprocessing

Data preprocessing was conducted using R (v4.4.1). Continuous cell migration features (RMC, aspect ratio, area, and adherence) were aggregated as mean values per patient (∼50 cells per patient per condition) across each experimental condition (stiffness, ECM, or drug) for each patient. To stabilize variance and minimize the influence of outliers, continuous variables were transformed with the BoxCox procedure (caret v7.0-1). This family of power transformations reduces skewness and approximates a normal distribution by identifying an optimal parameter (λ) for each feature. The transformed features were then centered to mean 0 and scaled to unit variance (Z score standardization). Categorical variables were one-hot or integer-encoded as appropriate.

#### Hierarchical clustering

For hierarchical clustering, features with ≤25% missing values were imputed using k-nearest-neighbors (k = 5; VIM v6.2.2). In sensitivity analyses, simple median imputation was also tested, which produced slightly different clusters but led to equivalent downstream results and interpretations. Clustering was performed using the Ward linkage method and Euclidean distance to define the FM and SM groups.

#### Survival analysis

To evaluate the prognostic value of the migratory phenotypes, progression-free survival (PFS) was utilized as the primary clinical endpoint. PFS was defined as the time interval (in months) from the date of initial surgical resection to the date of first documented radiographic disease recurrence or death from any cause. Survival probabilities were estimated using the Kaplan-Meier method and compared using log-rank tests. Survival analysis was conducted using survival (v3.6-4) and visualized using survminer (v0.5.0) in R.

### RNA-sequencing

#### RNA-seq

Eight primary tumor samples were collected at the time of surgical resection from glioblastoma patients treated at the University of Minnesota Medical Center between 2017 and 2019. Tumor samples were immediately placed in RNALater solution and flash frozen in liquid nitrogen and stored in –80 °C. RNA extraction and sequencing were performed at the University of Minnesota Genomics Center (UMGC, Minneapolis, MN). Total RNA was isolated with the RNAeasy Plus Universal Mini kit (Qiagen) and poly-A libraries were prepared using TruSeq-stranded mRNA (Illumina). Next-generation sequencing was performed on an Illumina HiSeq 2500 device in high output mode, generating 51 bp reads with ∼20 million paired reads per sample.

#### Preprocessing

Raw sequencing data was processed using the nf-core RNA-seq pipeline (v3.18.0) with default, reverse-stranded settings. Briefly, reads were checked for the presence of adapters and quality with FastQC (v0.12.1). Adapter sequences and low-quality bases were removed with Trim Galore! (v0.6.10) and Cutadapt (v4.9). The trimmed reads were mapped against the Ensembl human reference genome (GRCh38 release 113) with a spliced-aware aligner, STAR (v2.7.11b). Duplicate alignments were marked with Picard MarkDuplicates (v3.1.1). The output SAM files from STAR were sorted and indexed to BAM files using SAMtools (v1.21). Gene counts were generated using BAM-level quantification with Salmon (v1.10.3), and StringTie (v2.2.3) was used for transcript assembly and quantification.

### Mayo Clinic PDX RNA-seq

To supplement our primary patient RNA-seq and increase statistical power, we obtained RNA-seq data from ten Mayo Clinic Brain Tumor Patient-Derived Xenograft (PDX) lines (www.cbioportal.org) [37] with matching migration data from our previous study [10]. Migration data was processed as described above. Raw counts for Mayo Clinic Brain Tumor PDX samples were obtained via request from Mayo Clinic.

### Transcriptomic analysis

#### Differential expression analysis

Differential gene expression (DE) analysis was performed in R (v4.4.1) with DESeq2 (v1.44.0). For primary tumor samples, transcript abundance estimates from Salmon were converted to gene-level counts with tximport (v1.32.0), while raw counts were imported directly for PDX samples. The two count matrices were then merged for downstream analysis resulting in a total of 18 samples (8 primary and 10 Mayo PDX). Low-expressed genes were pre-filtered by only retaining genes with a count of at least 10 for at least half of the samples. To account for potential confounding effects, sample origin (primary vs. PDX) was included as a covariate in the design formula (∼origin + group).

The 18 patients were stratified into FM and SM groups using the combined RMC Z score across 0.7, 4.6, and 9.3 kPa substrates, with a Z score of 0 serving as the cutoff. Standard DE analysis between the FM and SM groups was performed (**Supplementary Table 3**). Normalization of counts for downstream analysis was performed by variance stabilized transformation (VST). Shrinkage of effect size (LFC estimates) for visualization and gene ranking was performed using the ashr method. Batch effects attributable to sample origin were corrected with limma (v3.60.5). Principal component analysis (PCA) was conducted on the batch-corrected VST-normalized counts to assess sample clustering using prcomp. Heatmaps were generated with ComplexHeatmap (v2.20.0) and volcano plots were generated with EnhancedVolcano (v1.20.0). Other plots were generated or modified using ggplot2 (v3.5.1).

### Pathway analysis

Over representation analysis (ORA) was performed using the R package clusterProfiler (v4.12.3) to identify significantly enriched (adjusted *P* value < 0.05) gene ontology (GO) pathways associated with migratory phenotypes. Human Entrez Gene identifiers were mapped using org.Hs.eg.db (v3.19.1), with the statistical background defined as all genes that passed expression filtering. Separate analyses were conducted for DEGs upregulated in the FM group (FDR < 0.05 and logFC > 0) and SM group (FDR < 0.05 and logFC < 0). Enrichment *P* values were calculated using a one-tailed hypergeometric test and adjusted for multiple comparisons using the Benjamini-Hochberg procedure (**Supplementary Table 4**). The results were visualized using enrichplot (v1.22.0) and ggplot2 (v3.5.1). Enrichment map networks were generated using emapplot to identify pathways with overlapping gene sets.

### Motor–clutch analysis

To quantify the relative expression of motor versus clutch genes, we defined two gene sets based on canonical genes in the motor–clutch model [26, 27]. The motor gene set included non-muscle myosin II heavy-chain isoforms MYH9, MYH10, and MYH14, which generate contractile forces that drive retrograde actin flow. The clutch gene set included the integrins ITGB1 and ITGA5, the talin isoforms TLN1 and TLN2, and vinculin (VCL), which couple the actin cytoskeleton to the extracellular matrix. Of the eight primary tumor samples with RNA-seq, five had corresponding blebbistatin migration data and were classified as blebbistatin-enhanced (n = 3) or blebbistatin-suppressed (n = 2) based on their response to myosin II inhibition. For each of these samples, a motor score and a clutch score were computed as the mean VST-normalized expression across the corresponding gene sets. Because VST values are on a log-like scale, a motor–clutch logFC ratio was obtained by subtracting the clutch score from the motor score.

We also derived DEGs and enriched pathways for the blebbistatin-enhanced and -suppressed groups as described above (**Supplementary Table 5, Supplementary Table 6**). To assess whether the results could be replicated in an unsupervised manner, we only used motor and clutch genes found in the DEGs. We manually identified motor and clutch genes among the DEGs apriori and computed motor/clutch ratios. Due to the limited sample size (n = 5), formal statistical comparisons of motor–clutch differences were not performed.

### Migration gene signature

#### TCGA and CGGA Transcriptomics

We obtained RNA-seq RSEM values from The Cancer Genome Atlas (TCGA) glioblastoma dataset (www.cbioportal.org, Cell 2013 [35], n = 144 when IDH1-mutant excluded) and raw counts from the Chinese Glioma Genome Atlas (CGGA) (www.cgga.org.cn [38], mRNAseq_325, mRNAseq_693, n = 298 when IDH1-mutant tumors excluded). For these RNA-seq datasets, raw count matrices were imported via DESeqDataSetFromMatrix, with RSEM values rounded to the nearest whole number. Low-expressed genes were pre-filtered by retaining only genes with ≥10 counts in at least 20 samples (TCGA) or 200 samples (CGGA), followed by normalization via variance stabilized transformation (VST). Additionally, we obtained pre-normalized expression data from the TCGA glioblastoma Firehose microarray dataset (www.cbioportal.org, Firehose Legacy, n = 525). Notably, IDH1-mutational status was not available for the microarray cohort, and its patient population contains partial overlap with the TCGA RNA-seq cohort.

### Signature derivation

To derive a migration signature with prognostic value, we iteratively pruned the 141 DEGs between the FM and SM groups using the TCGA RNA-seq dataset (n = 144, IDH1-mutant excluded) as a discovery cohort. At each iteration, we removed the gene whose exclusion maximized Harrell’s concordance index (c-index) for progression-free survival (PFS). The c-index measures predictive discrimination in survival analysis and is the fraction of all usable patient pairs in which the subject with the higher predicted risk experiences the event sooner (0.5 = chance, 1 = perfect). After each pruning step we also computed log-rank *P* value (Kaplan-Meier, KM), Likelihood ratio test *P* value (Cox), time-dependent area under the receiver operating curve (AUC) at median survival, and integrated Brier score (IBS) to further assess model performance.

Rather than strictly selecting the model with the absolute maximum c-index, which may lead to overfitting on a small number of genes, we applied a tolerance-based selection criterion. We identified the maximum c-index achieved during pruning and selected all candidate gene sets that performed within 0.01 of this peak. From these candidate sets, we selected the largest gene set, yielding a robust 13-gene migration signature (**Supplementary Table 7**). To assess the specificity of the signature, we performed a permutation test with 10,000 bootstrap iterations. In each iteration, a random 13-gene set was sampled from the DEGs, and its prognostic performance was compared to the migration signature.

### Prospective *in vitro* validation

For prospective validation of the migration signature, we calculated migration scores for the complete cohort of Mayo PDX samples with available RNA-seq as described previously. For each patient, we computed the migration score using single-sample gene set enrichment scoring with singscore (v1.24.0), defined as the difference between independently computed rank-based FM and SM gene set scores (FM − SM). Based on these scores, we stratified samples and prospectively selected eight PDX lines representing distinct predicted phenotypes: four predicted as FM and four predicted as SM. These eight previously uncharacterized lines were then subjected to 2D migration assays on laminin-coated PA gels with stiffness levels mimicking brain tissue (4.6 kPa) in a blinded manner. Upon unblinding, RMC was quantified to determine whether migration scores correlated with observed motility, as described above.

### Prospective *in vivo* validation

We calculated migration scores for each PDX sample using single-sample gene-set enrichment (singscore, v1.24.0). We then examined the relationship between these migration scores and *in vivo* invasiveness, using histologically defined invasion stages (I–IV) from the Mayo Clinic [37]. Briefly, invasion was quantified histologically on coronal brain sections by immunostaining for human-specific markers (e.g., Lamin A/C) to identify infiltrating tumor cells and assign an ordinal invasion score (I–IV), ranging from unilateral, well-demarcated tumors to diffuse bilateral infiltration with midline crossing, based on the extent and pattern of parenchymal spread.

### Survival analysis

For each patient, migration signature scores were computed using single-sample gene-set enrichment (singscore, v1.24.0). Patients were stratified into high-(FM) and low-score (SM) groups based on cohort-specific median scores, and survival analysis was performed. We validated the migration signature in two independent cohorts, the larger CGGA cohort for overall survival (OS) and TCGA RNA-seq and microarray cohorts for PFS and OS. Kaplan-Meier curves were compared using log-rank tests (survival v3.6-4).

To determine whether the migration signature serves as an independent prognostic indicator when controlling for established clinical and molecular confounders, we performed multivariate Cox proportional hazards regression. The covariate adjustments were tailored to the available clinical data for each cohort. The TCGA analysis adjusted for molecular subtype (G-CIMP, Neural, Proneural, and Mesenchymal), MGMT promoter methylation status, sex, and age.The CGGA analysis adjusted for tumor type (primary, secondary, recurrent), MGMT promoter methylation status, sex, and age. All survival analyses were conducted using survival (v3.6-4) and visualized using survminer (v0.5.0) in R.

### Orthogonality analysis

To evaluate the orthogonality of the migration signature relative to existing glioblastoma subtypes, we benchmarked our 13-gene signature against four published gene sets: the standard molecular subtypes (Proneural, Classical, Mesenchymal) from Wang et al. (2017) [5]; developmental and injury response programs from Richards et al. (2021) [9]; metabolic states (Glycolytic/Plurimetabolic, Proliferative/Progenitor, Mitochondrial, Neuronal) from Garofano et al. (2021) [7]; and the lineage subtypes (Type I, Type II; renamed Neural Stem and OPC respectively) from Wang Z. et al. (2020) [8]. Gene set overlap was quantified using the Jaccard index. For each patient, migration signature scores were computed using single-sample gene-set enrichment (singscore, v1.24.0). The association between signatures was quantified using Kendall’s rank correlation coefficient (tau). This metric was selected because it measures the probability of concordant patient ordering rather than linear value association, making it robust to diverse gene signatures. *P* values for pairwise correlations were adjusted for multiple hypothesis testing using Bonferroni correction. To further determine if the migration signature was correlated with any existing glioblastoma subtypes, we stratified TCGA RNA-seq cohort patients (n=144, IDH1-mutant excluded) according to subtype assignments from each publication. Patients were assigned to the subtype yielding the maximal single-sample enrichment score among the signatures defined in each respective study. We compared the distribution of migration scores within each subtype against the cohort-wide median using Wilcoxon rank-sum tests.

### Single-nucleus RNA-sequencing

#### Patient samples

Multi-regional sampling was performed under an institutional review board–approved protocol (IRB120345X), with informed consent obtained from all patients and in accordance with the Declaration of Helsinki. Using Medtronic stereotactic guidance, neurosurgeons secured targeted specimens from three predefined anatomical compartments: the tumor core (non-enhancing), the contrast-enhancing tumor edge, and adjacent normal cortex (non-enhancing tissue beyond the tumor margin). To minimize brain shift, biopsies were obtained early during resection. Each specimen underwent immediate neuropathologic review to confirm that the sampled tissue corresponded to its intended spatial region, ensuring accurate assignment of core, edge, and normal compartments for downstream analyses. Tumor specimens were flash-frozen in liquid nitrogen immediately after surgical resection and stored at −80 °C until processing for 10x Genomics Multiome analysis.

### 10x Multiome

Samples were thawed and nuclei were isolated as previously described [50]. Isolated nuclei were processed using the 10x Genomics Chromium Single Cell Multiome ATAC + Gene Expression assay according to the manufacturer’s protocol, targeting recovery of 5,000–10,000 nuclei per sample. Library quality was assessed on an Agilent 4200 TapeStation using High Sensitivity ScreenTape, and library concentrations were quantified by real-time PCR. Libraries were sequenced on an Illumina NextSeq 2000 platform. Raw sequencing data were processed using Cell Ranger ARC (10x Genomics) and aligned to the human GRCh38 reference genome [40].

### CopyKAT

To identify malignant cells, we inferred large-scale copy number alterations using CopyKAT (tl.copykat) implemented in infercnvpy (v0.6.1) which classifies aneuploid tumor cells apart from diploid normal cells. Analyses were performed on raw UMI counts and CNV profiles with a smoothing window of 25 genes. Cells called as aneuploid by CopyKAT largely matched expression-based tumor annotations. CopyKAT was run on each sample individually before integration and inference was performed without a reference since CopyKAT uses an unsupervised Bayesian approach that identifies diploid and aneuploid populations directly from the data.

### snRNA-seq analysis

All analyses of single-nucleus RNA-sequencing (snRNA-seq) data was performed in Python (v3.13.7) using Scanpy (v1.11.4) following standard best practices [51]. In brief, cells with low total counts or excessively high mitochondrial RNA fraction were excluded, and genes expressed in only a few cells were filtered. For integration of multiple samples, we applied Harmony with default parameters to correct for batch effects while preserving biological variation. Counts were library-size normalized (sc.pp.normalize_total) and log-transformed (sc.pp.log1p). Highly variable genes were selected (sc.pp.highly_variable_genes), data were scaled to unit variance, principal components (PCs) were computed (sc.tl.pca), and a neighborhood graph was built (sc.pp.neighbors). Two-dimensional embeddings were generated with UMAP (sc.tl.umap) for visualization. Clustering was performed using Leiden on the kNN graph. Manual cell type annotation was performed using a combination of established cell type markers and automated reference-based annotation using CellTypist (v1.4.0). Cells showing concurrent expression of marker genes from more than one cell type were flagged as potential doublets and filtered as appropriate. In addition, computational doublet detection was performed in Scanpy using Scrublet (v0.2.3).

To quantify a per-cell migration score we used our 13-gene migration signature and scored each cell using Scanpy gene-set scoring (sc.tl.score_genes). This function computes the average expression of the query gene set minus the average expression of a matched reference set of control genes sampled from the same expression bins. We computed two scores, one for the up-regulated list (Fast genes) and one for the down-regulated list (Slow genes) and defined the migration score as up minus down. Migration scores were standardized across cells (Z score) and used to categorize cells as slow (z < 0) or fast (z ≥ 0). For each region we calculated the fraction of cells in each state (SM or FM), the log2-fold change of FM to SM cells, and the average migration score.

### Validation scRNA-seq

To validate the spatial relevance of our migration signature, we utilized a 5-ALA-guided, region-annotated glioblastoma single-cell RNA sequencing (scRNA-seq) dataset from Mossi Albiach et al. 2025, publicly available on CZ CELLxGENE Discover [41]. This sample was obtained from a right temporal lobectomy, a surgical procedure that enabled systematic multi-region sampling extending significantly beyond the standard resection margin. Tissue biopsies were harvested along a radial trajectory from the tumor mass into macroscopically normal neocortex and annotated via intraoperative 5-ALA fluorescence to minimize brain shift bias. The anatomical labels included macroscopically normal cortex (C), periphery (P), tumor regions with low (LF) or high (HF) fluorescence, and necrotic core (NC). We used the authors’ sampling scheme and region labels as provided. Raw count matrices and accompanying metadata were downloaded directly from CELLxGENE.

### Magnetic Resonance Elastography (MRE)

To characterize the stiffness gradients of glioblastoma tumors *in vivo*, we reanalyzed publicly available MRE data from 9 glioblastoma patients. Imaging data was obtained from Svensson et al. (2021) [17] available via Zenodo (record 4926005). The dataset contains co-registered multimodal MR images, corresponding segmentations, and quantitative MRE stiffness maps for patients with histologically confirmed glioblastoma. All images were processed in a common Montreal Neurological Institute (MNI) space to enable spatially consistent radial sampling across patients. All analyses were conducted in Python (v3.10)

For each subject, MRI-derived segmentation masks defined four distinct regions: necrotic core (index 0), contrast-enhancing tumor (1), peritumoral edema (2), and normal-appearing brain/cortex (3). All computations were restricted to voxels within the brain mask. For elastography, we used the magnitude of the complex shear modulus (Gabs, |G*|). To express tissue mechanics in a form more comparable to our 2D migration assays, we converted |G*| to an approximation of the isotropic Young’s modulus using the relation E = 3|G*|, assuming tissue incompressibility (Poisson’s ratio ≈ 0.5). This conversion is standard in the field and well-suited to brain tissue, which is highly hydrated and exhibits nearly incompressible behavior under small deformations. We applied 3D Gaussian smoothing (σ = 2 voxels) to suppress voxel-level noise.

To characterize how stiffness changes from the center of the tumor to the surrounding brain, for every subject, we identified the tumor center as the point deepest inside the necrotic/contrast-enhancing compartment using a distance-transform-based approach (voxel with the maximal distance to the necrotic core boundary). Distances from this center to all intracranial voxels were computed in 3D and binned into concentric shells from 0 to 100 mm in 2-mm steps. For each shell we calculated (i) the mean MRE-derived stiffness (with 5% trimming to limit the influence of outliers); (ii) the corresponding variance; (iii) the average tissue index (0–3) as a proxy for the dominant region at that radius; and (iv) the proportion of voxels belonging to each of the four regions. One subject (Patient F) was excluded after analysis due to its atypical stiffness profile (extremely stiff tumor core, >8 kPa).

### Statistics and reproducibility

All statistical analyses were performed in R (v4.4.1) with data manipulation and visualization in the tidyverse ecosystem (v2.0.0). Mixed-effects modeling was conducted using lme4 (v1.1-35), with Type-III Analysis of Variance (ANOVA) tables and Kenward-Roger degrees of freedom approximations computed via lmerTest (v3.1-3). Post-hoc marginal means and pairwise comparisons were estimated using emmeans (v1.10.0). Model assumptions (normality of residuals, homoscedasticity) were verified using performance (v0.14.0) and insight (v1.3.1).

To ensure rigorous statistical inference and avoid pseudoreplication, biological replicates were defined as individual patients, while single cells were treated as subsamples (technical replicates) nested within patient. While data were collected at the single-cell level, to prevent inflation of statistical power, primary hypothesis testing was conducted on patient-level aggregated means. Single cells were modeled as repeated measures using appropriate random effects structures. When patients were measured on multiple substrates, these measurements were also treated as repeated measures within the same patient. This approach ensured that *P* values reflected the variability between biological donors rather than intra-patient technical noise.

Consequently, we used hierarchical linear mixed-effects models (LMMs) to model patient means using lme4, incorporating random intercepts for each patient to account for the repeated-measures design (where the same patient is measured across multiple stiffnesses or substrates).

For migration metrics, we used a hierarchy of linear mixed-effects models with varying random effects structures: random intercepts for patient effects, with or without random slopes for patient × substrate interactions. Models were fit to both cell-level data and patient-aggregated means. Models were evaluated using AIC/BIC and marginal/conditional R^2^.

Stiffness was modeled as a fixed effect (log10 transformed to linearize the exponential range of matrix elasticity). To determine whether cellular mechanosensitivity followed a linear or biphasic response across the stiffness range (0.7–100 kPa), we compared two regression frameworks: (1) a log-linear model assuming a continuous, monotonic response proportional to log10(stiffness) and (2) a biphasic (piecewise linear) model assuming a change in slope at particular knots defined a priori (4.6 kPa for RMC or the geometric mean of 4.6 and 9.3 kPa for aspect ratio). For biphasic models, the significance of the shift in the stiffness response was evaluated by testing the difference between the pre-and post-breakpoint slopes using a Wald t-test. For group comparisons (FM vs. SM patients), we assessed statistical significance by the group main effect. Significance of fixed effects was determined using Type III ANOVA with Kenward-Roger degrees of freedom approximations. Post-hoc testing was performed to compare differences between groups at specific stiffness levels and differences across stiffness levels with Tukey’s adjustment for multiple comparisons.

For experiments involving distinct ECM coatings (laminin, collagen, Matrigel, fibronectin), substrate ECM was modeled as a fixed categorical effect. For group comparisons (FM vs. SM patients), we assessed statistical significance based on the group main effect. A significant interaction indicated that the two groups exhibited distinct responses to ECM. Significance of fixed effects was determined using Type III ANOVA with Kenward-Roger degrees of freedom approximations. Post-hoc testing was performed to compare differences between groups on specific ECMs and differences across ECMs with Tukey’s adjustment for multiple comparisons.

Due to the limited sample size in drug perturbation experiments (latrunculin A, n = 6; blebbistatin, n = 17) preventing robust parametric modeling, comparisons between treated and untreated conditions were performed using non-parametric two-sided Wilcoxon rank-sum tests. For completeness, all mixed-effects models were also fit to the raw cell-level data (nesting cells within patients via random effects). In all cases, fixed-effect conclusions derived from cell-level models were consistent with those from the primary patient-mean analysis. Statistical significance was defined as p < 0.05.

## Data availability

RNA-seq data generated in this study will be deposited in the NCBI Gene Expression Omnibus (GEO) upon publication. Publicly available datasets used in this study include: The Cancer Genome Atlas (TCGA) glioblastoma RNA-seq dataset, available via cBioPortal under the study ID gbm_tcga_pub2013; The TCGA glioblastoma microarray dataset, available via cBioPortal under the study name: Firehose Legacy; the Chinese Glioma Genome Atlas (CGGA) RNA-seq datasets, available via the CGGA Portal under dataset IDs mRNA-seq_325; the Mayo Clinic Brain Tumor Patient-Derived Xenograft (PDX) RNA-seq dataset, available via cBioPortal under study ID gbm_mayo_pdx_sarkaria_2019; and the 5-ALA guided region-annotated glioblastoma scRNA-seq dataset, available via the CZ CELLxGENE Discover repository under dataset SL040 [41, 52]. Counts for Mayo Clinic Brain Tumor PDX samples are not publicly available and were obtained via request from Mayo Clinic. Magnetic Resonance Elastography (MRE) data are available via Zenodo (record 4926005) [17]. Source data are provided with this paper.

## Code availability

All code will be publicly available in a GitHub repository and archived at Zenodo upon publication.

## Acknowledgements

This work was supported by the NIH (grant nos. U54 CA210190, P01CA254849, U54 CA268069, and R01-CA-172986 to D.J.O., grant nos. 5P20GM109035-10 to J.H). We thank Dr. Matthew Hunt for performing the surgeries to obtain the clinical specimens used in this study. Z.W. is a DDBrown Awardee of the Life Sciences Research Foundation. RNA-seq analysis was supported by Juan E. Abrahante Lloréns at the University of Minnesota Genomics Center. Figures were created using BioRender and Adobe Illustrator. Computational resources were provided in part by Brown University’s high-performance computing cluster, OSCAR. Above all, we are deeply grateful to the patients and their families, whose generosity and willingness to participate made this research possible.

## Author Information

**These authors jointly supervised this work:**

Clark C. Chen and David J. Odde.

**Contributions:**

J.H. and M.M. performed the experiments. A.W.N. and J.H. performed the sequencing and bioinformatic analyses. A.W.N. and J.H. performed statistical analyses. C.C.C., J.H., and D.O., designed the experimental set-up and conceived the study. Z.W., T.H.L., Y.X., and L.C contributed to 10x Multiome data generation and A.W.N., Z.W., and B.T. performed analyses. B.R. supervised 10x Multiome data generation. A.G., and C.C.C. contributed patient samples and clinical resources. Z.G. provided institutional support and resources. S.I.D. provided statistical oversight and manuscript review. C.C.C. and D.O. supervised the research and obtained funding. A.W.N. and J.H. wrote the manuscript. All authors proofread and approved the final paper.

## Supplementary information

**Supplementary Tables:** Supplementary Tables 1–7.

Supplementary Table 1: Patient clinical characteristics.

Supplementary Table 2: CMS parameters.

Supplementary Table 3: Differential expression analysis (FM vs. SM).

Supplementary Table 4: Pathway enrichment analysis (FM vs. SM).

Supplementary Table 5: Differential expression analysis (blebbistatin-enhanced vs. -suppressed).

Supplementary Table 6: Pathway enrichment analysis (blebbistatin-enhanced vs. -suppressed).

Supplementary Table 7: The 13-gene migration signature.

**Supplementary Figures:** Supplementary Figures 1–7.

## Source data

Source Data is available for Figs. 1–5, Extended Data Figs. 1–10, Supplementary Figs. 1–7.

---

**Extended Data Fig. 1 (related to Fig. 1):**
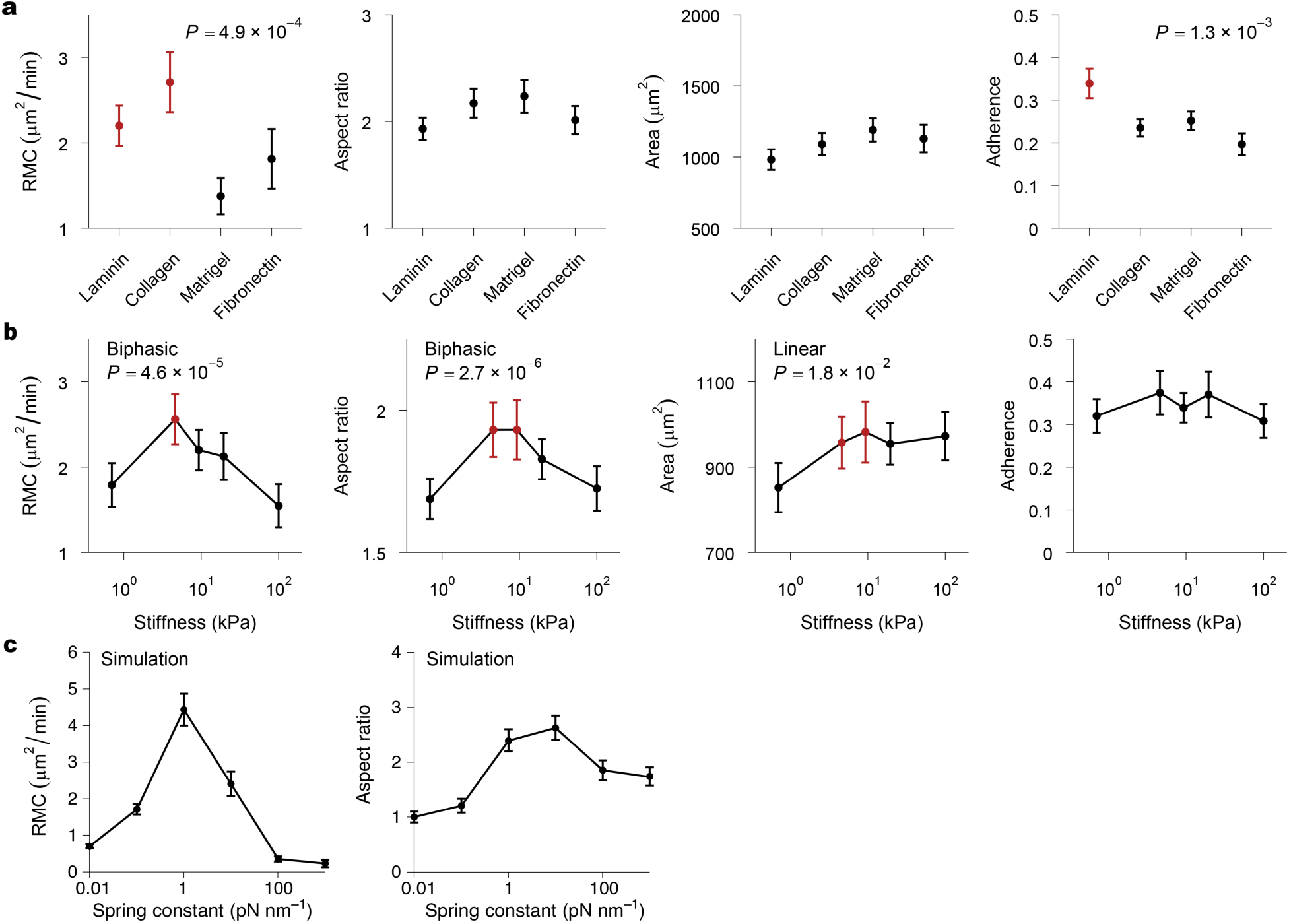
ECM ligand and stiffness screening identify optimal conditions for glioblastoma migration. **a, b,** Screening of ECM ligands (**a**) and stiffness levels (**b**) to identify the optimal substrate for migration. Quantification of random motility coefficient (RMC), aspect ratio, area, and fractional adherence reveal laminin and collagen promote significantly higher RMC (*P* = 4.9×10^−4^) and laminin promotes greater substrate adherence (*P* = 1.3×10^−3^) compared with other ligands (Type III ANOVA, linear mixed-effects models). Thus, laminin was selected as the ligand for experiments with varying stiffness. RMC (*P* = 4.6×10^−5^) and aspect ratio (*P* = 2.7×10^−6^) exhibit a biphasic dependence on stiffness, with maxima at intermediate, brain-like stiffness (∼4.6–9.3 kPa) (Wald t-test on change in slope). Data are mean ± s.e.m. Red points indicate significantly higher post hoc comparisons (see **Supplementary** Fig. 1). **c,** Computational motor–clutch model simulations for RMC and aspect ratio as a function of substrate stiffness (data from [15]).

**Extended Data Fig. 2 (related to Fig. 1):**
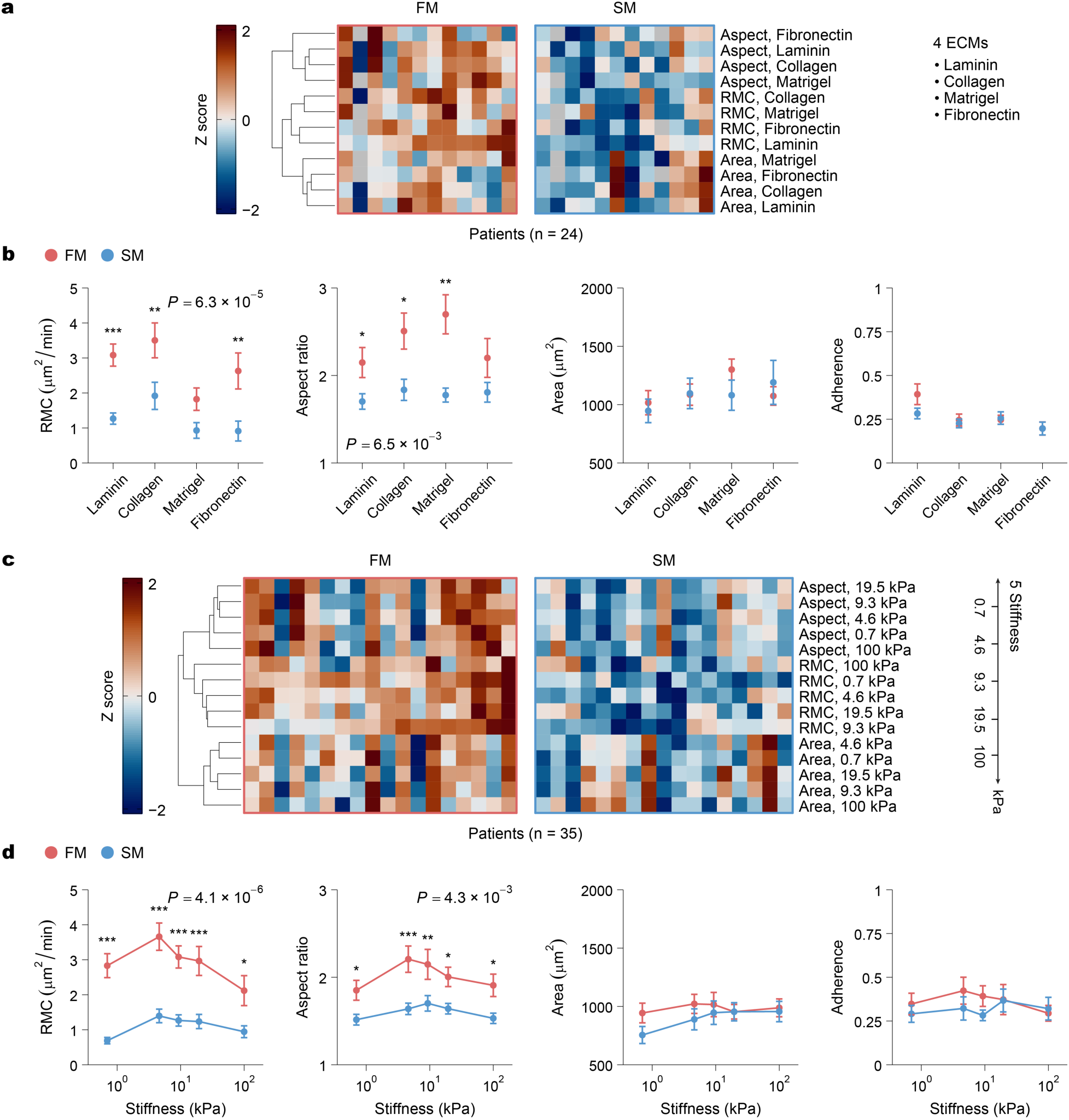
Fast-Migrating (FM) glioblastoma cells exhibit enhanced sensitivity to ECM composition and substrate stiffness. **a**, Heatmap of Z scored migration features (random motility coefficient (RMC), aspect ratio, area, and adherence) across 9.3 kPa substrates coated with laminin, collagen, Matrigel, or fibronectin for n = 24 patient samples. Patients (columns) are ordered as in Fig. 1b and grouped into FM and SM phenotypes. Hierarchical clustering (Euclidean distance, Ward linkage) of migration metrics (rows) separates area, RMC, and aspect ratio across ECM conditions. **b**, Migration metrics from **a** stratified by FM (red) and SM (blue) phenotypes. *P* values indicate the significance of the FM versus SM group effect (Type III ANOVA), and asterisks denote significant Tukey-adjusted post hoc comparisons between FM and SM within each ECM condition. **c**, Heatmap (as in **a**) of migration features across laminin-coated substrates spanning stiffnesses of 0.7, 4.6, 9.3, 19.5, and 100 kPa for n = 35 patient samples. Hierarchical clustering (Euclidean distance, Ward linkage) of migration metrics (rows) separates area, RMC, and aspect ratio across substrate stiffness. **d**, Migration metrics from **c** stratified by FM (red) and SM (blue) phenotypes. *P* values indicate the significance of the FM versus SM group effect (Type III ANOVA), and asterisks denote significant Tukey-adjusted post hoc comparisons between FM and SM at each stiffness level. Significance: ns, *P* ≥ 0.05; *, *P* < 0.05; **, *P* < 0.01; ***, *P* < 0.001.

**Extended Data Fig. 3 (related to Fig. 1):**
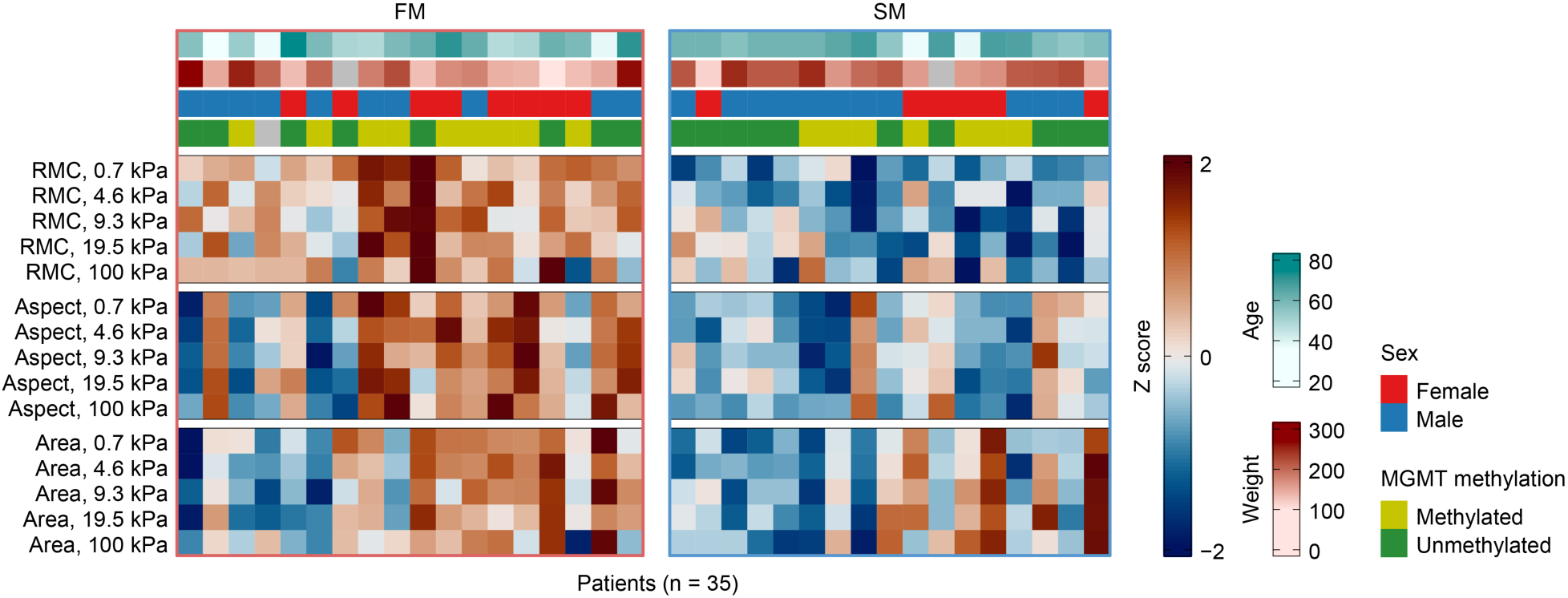
Association of clinical variables with the migratory phenotypes. Heatmap of migration features (random motility coefficient (RMC), aspect ratio, and area) on laminin-coated 0.7, 4.6, 9.3, 19.5, and 100 kPa substrates. Columns represent individual patients (n = 35), clustered into FM (red) and SM (blue) migratory phenotypes as defined in Fig. 1b. Clinical variables (top) include age, weight, sex, and MGMT promoter methylation (**Supplementary Table 1**). Migratory phenotype was not associated with any of these clinical variables (*P* > 0.05, two-sided Chi-square test for categorical variables, two-sided Wilcoxon rank-sum test for continuous variables).

**Extended Data Fig. 4 (related to Fig. 2):**
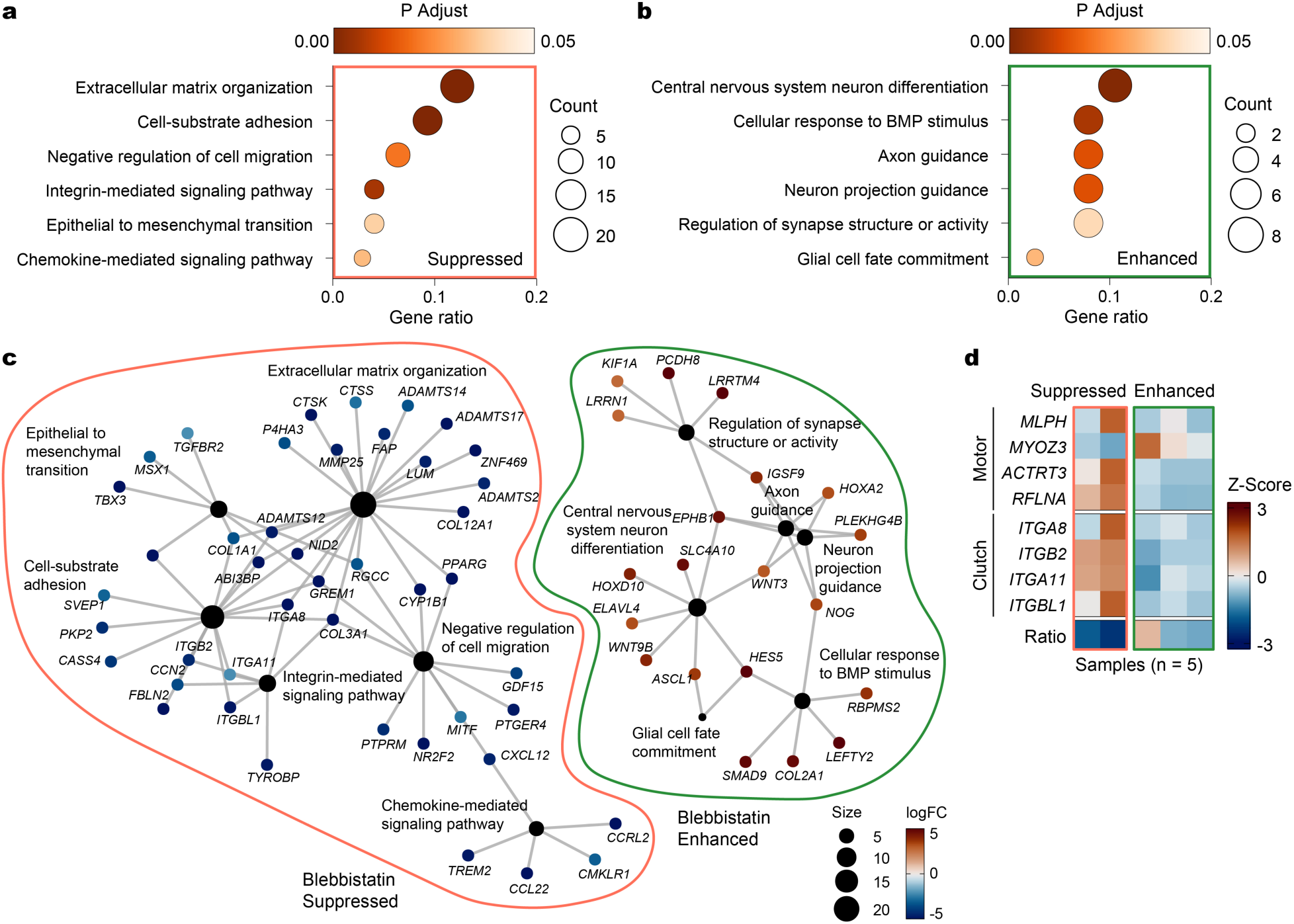
Transcriptional pathways distinguish blebbistatin-enhanced and blebbistatin-suppressed motor–clutch states. **a, b,** Dot plot of pathways enriched in blebbistatin-suppressed DEGs (**a**, orange) and blebbistatin-enhanced DEGs (**b**, green). Gene Ratio denotes fractions of genes annotated to a given GO term relative to the total genes in that term. Dot size represents gene count. Color indicates statistical significance (adjusted *P* value). **c,** Enrichment map network visualizing links between GO pathways (black nodes) and genes (colored nodes). Pathway node size is proportional to the number of mapped genes, while gene nodes are colored by log2-fold change (logFC). The network highlights an ECM and migration-regulatory module specific to blebbistatin-suppressed DEGs (orange) and a neuronal differentiation and guidance module specific to blebbistatin-enhanced DEGs (green). **d,** Heatmap showing row-scaled Z scores of selected motor and clutch DEGs (**see Fig. 2j,k**) in blebbistatin-suppressed versus blebbistatin-enhanced patients (n = 5).

**Extended Data Fig. 5 (related to Fig. 3):**
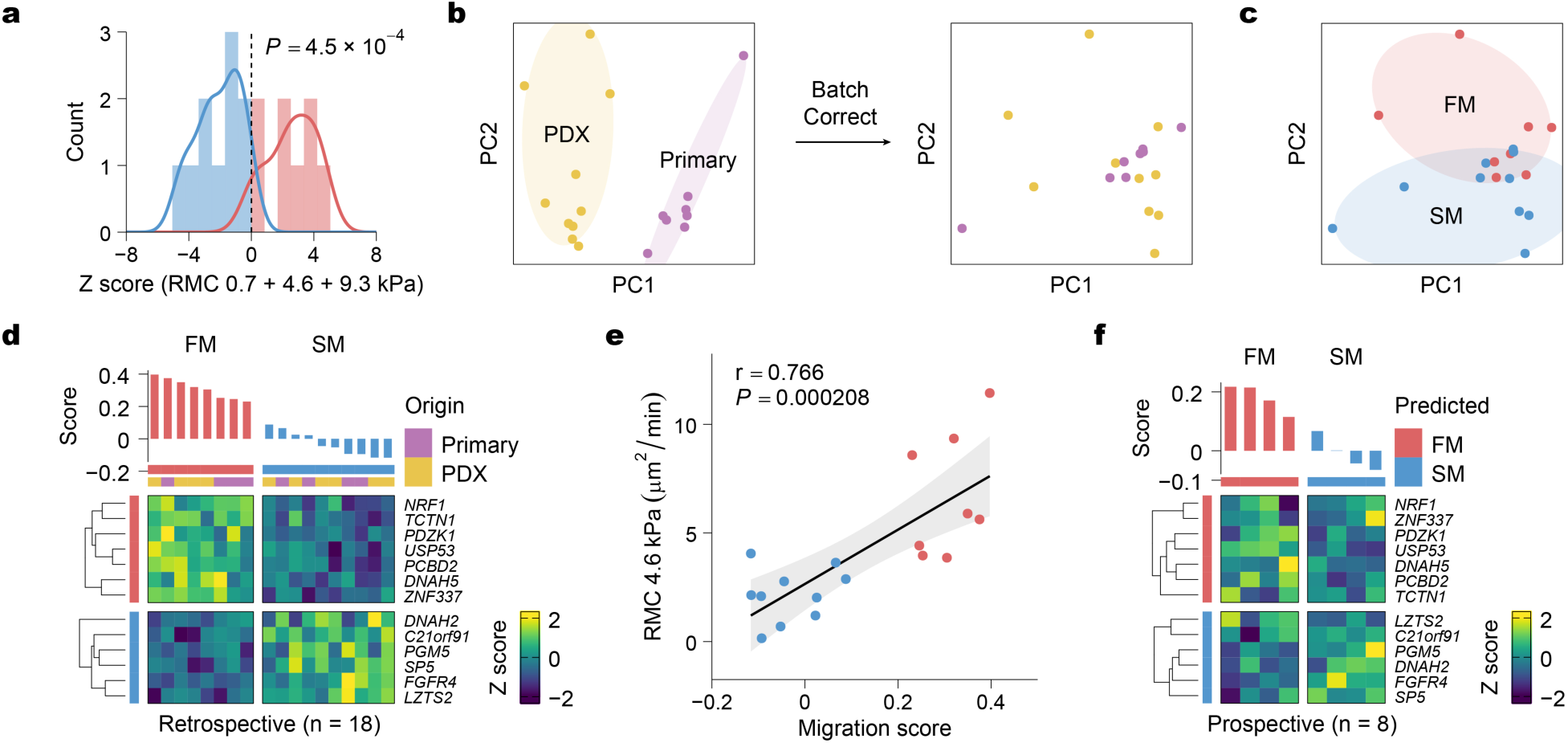
Discovery and validation of the migration gene signature. **a,** Histogram showing stratification of patients into Fast-Migrating (FM) and Slow-Migrating (SM) groups for differential expression analysis (Fig. 3c) using a combined RMC Z score of 0 as the cutoff (dashed line) across 0.7, 4.6, and 9.3 kPa substrates. The FM (red) and SM (blue) groups display significantly different distributions (*P* = 4.5×10^−4^, two-sided Wilcoxon rank-sum test). **b,** Principal component analysis (PCA) of RNA-seq data from Mayo PDX tumors (yellow, n = 10) and primary tumors (magenta, n = 8) before (left) and after (right) limma batch correction. **c,** PCA of the batch-corrected bulk RNA-seq data, showing separation into FM (red) and SM (blue) groups. **d,** Retrospective application of the migration signature to the original 18 samples. The distribution of migration scores (top) and heatmap of the 13-gene migration signature (bottom) demonstrates the migration signature accurately predicts migration in 100% of the samples. **e,** Retrospective validation of the migration signature to predict *in vitro* cell motility (random motility coefficient, RMC) on 4.6 kPa substrates (Pearson r = 0.766, *P* = 0.000208). **f,** Prospective validation of the signature in a previously uncharacterized cohort of Mayo PDX samples (n = 8). The distribution of migration scores (top) and heatmap of the 13-gene migration signature (bottom) shows the predicted migration for each of the 8 prospective samples.

**Extended Data Fig. 6 (related to Fig. 3):**
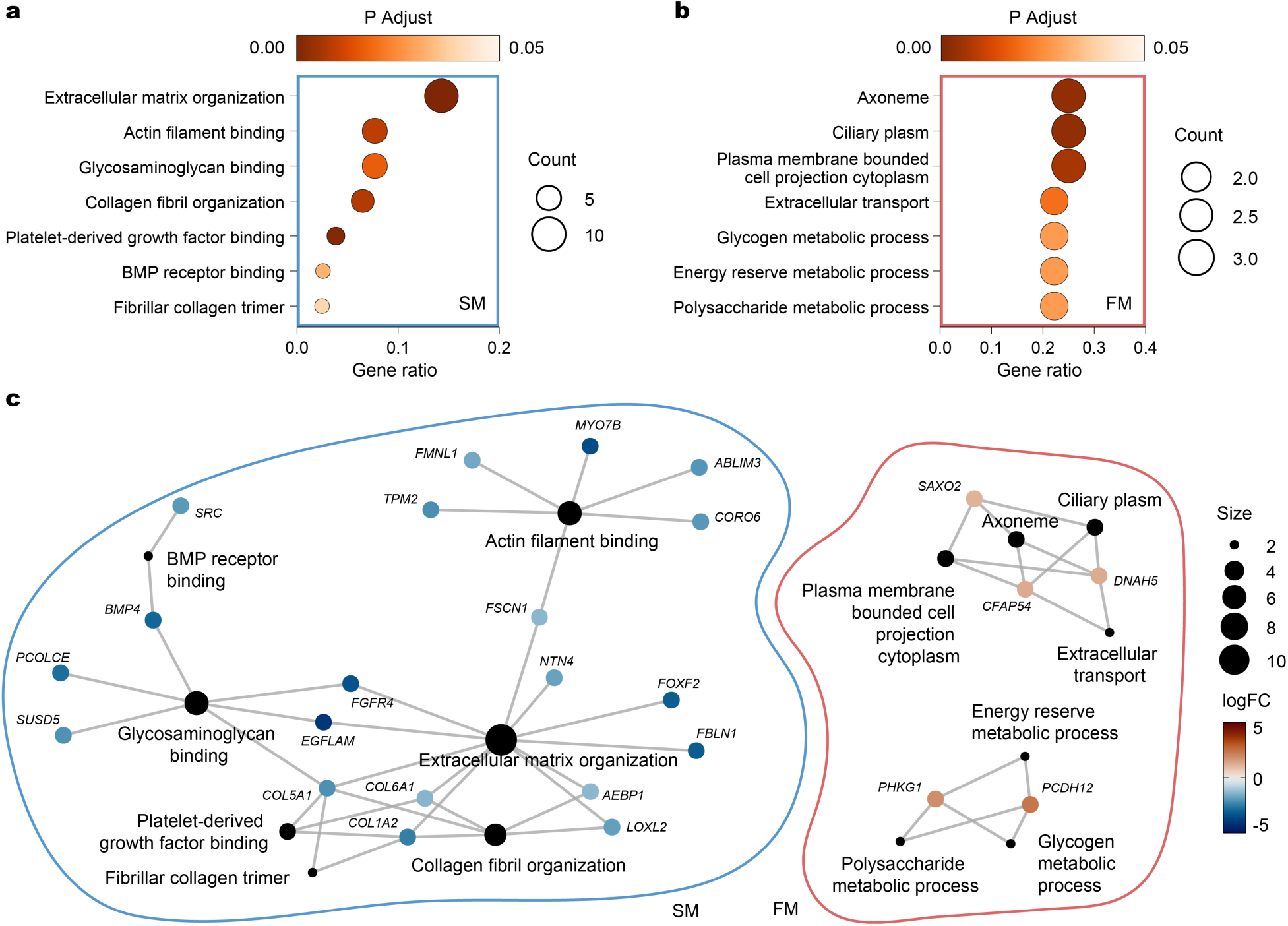
Transcriptional pathways distinguish FM and SM phenotypes. **a, b,** Dot plot enrichment analysis of DEGs upregulated in the Slow-Migrating (SM) (**a**, blue) and Fast-Migrating (FM) phenotypes (**b**, red). Gene Ratio denotes fractions of genes annotated to a given GO term relative to the total genes in that term. Dot size represents gene count. Color indicates statistical significance (adjusted *P* value). The SM phenotype **(a)** is enriched for ECM organization, collagen fibril organization, and BMP receptor binding while the FM phenotype **(b)** is enriched for ciliary plasm, plasma-membrane-bounded cell-projection cytoplasm, and glycogen metabolism process. **c,** Enrichment map network visualizing links between GO pathways (black nodes) and genes (colored nodes) shown in **(a)** and **(b)**. Pathway node size is proportional to the number of mapped genes, while gene nodes are colored by log2-fold change (logFC). The network highlights an ECM/adhesion module specific to SM DEGs (blue) and a ciliary/axonemal and glycogen-metabolism module specific to FM DEGs (red).

**Extended Data Fig. 7 (related to Fig. 3):**
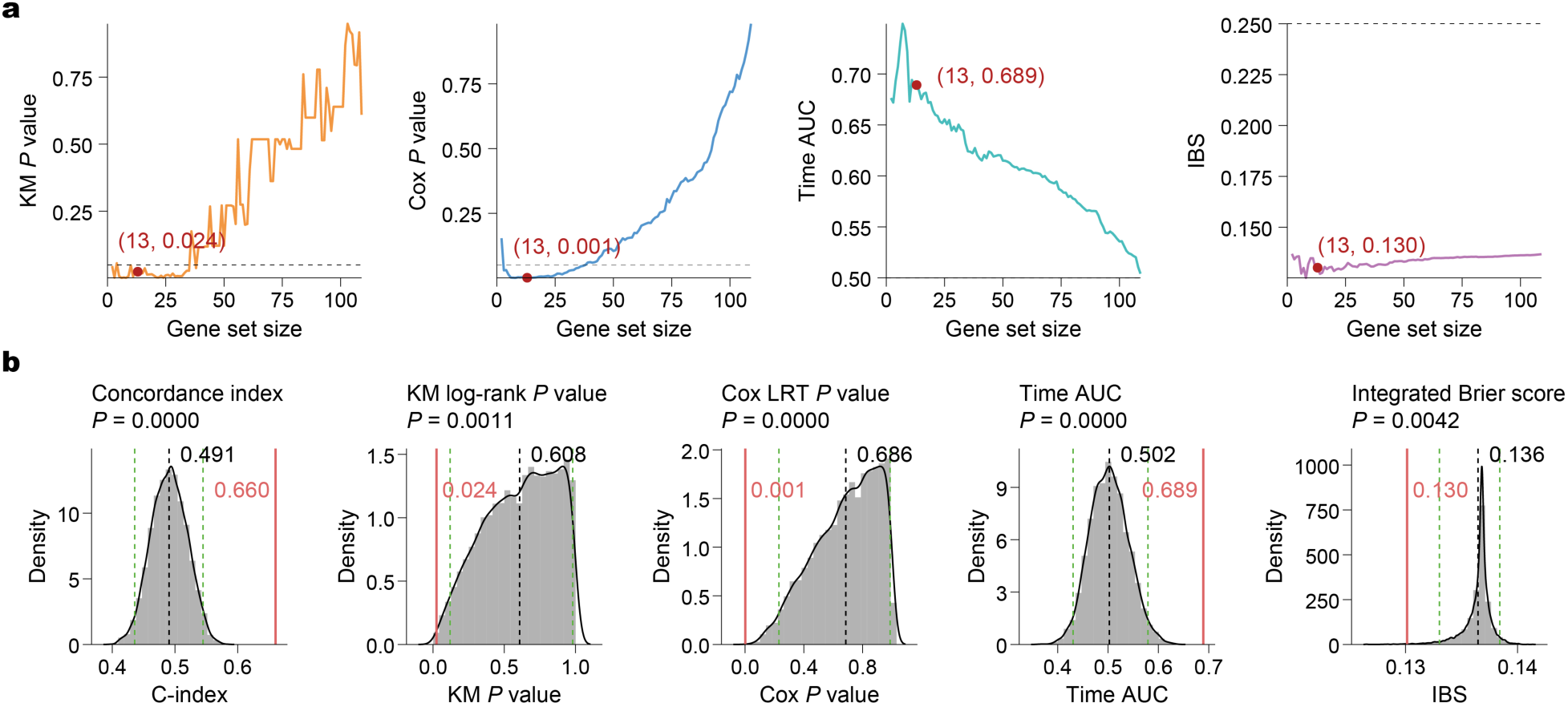
Optimization and validation of the migration gene signature. **a,** Stepwise optimization of the migration gene signature using the TCGA RNA-seq cohort (n = 144, IDH1-mutant excluded). Starting from all DEGs, genes were iteratively removed to maximize Harrell’s concordance index (see Fig. 3d). Plots display other performance metrics as a function of gene-set size including Kaplan-Meier (KM) log-rank *P* value (orange), Cox likelihood-ratio (LRT) test *P* value (blue), time-dependent area under the receiver operating characteristic curve (Time AUC, teal), and integrated Brier score (IBS, purple). Dashed horizontal lines denote null expectations or significance thresholds. The red dot marks the optimal, 13-gene subset. **b,** Permutation testing comparing the performance of the final 13-gene signature (solid red lines) to 10,000 randomly sampled gene sets from the pool of DEGs of identical size (gray histograms). Metrics evaluated include c-index, KM log-rank *P* value, Cox LRT *P* value, time-dependent AUC, and IBS. Blacked dashed lines mark the mean of the null distribution; green dashed lines mark the 95% confidence intervals. One-sided empirical *P* values were calculated as the proportion of random samples that performed as well as the observed signature. Across all five metrics the migration signature falls in the extreme tails indicating significantly better discrimination, calibration, and risk stratification.

**Extended Data Fig. 8 (related to Fig. 4):**
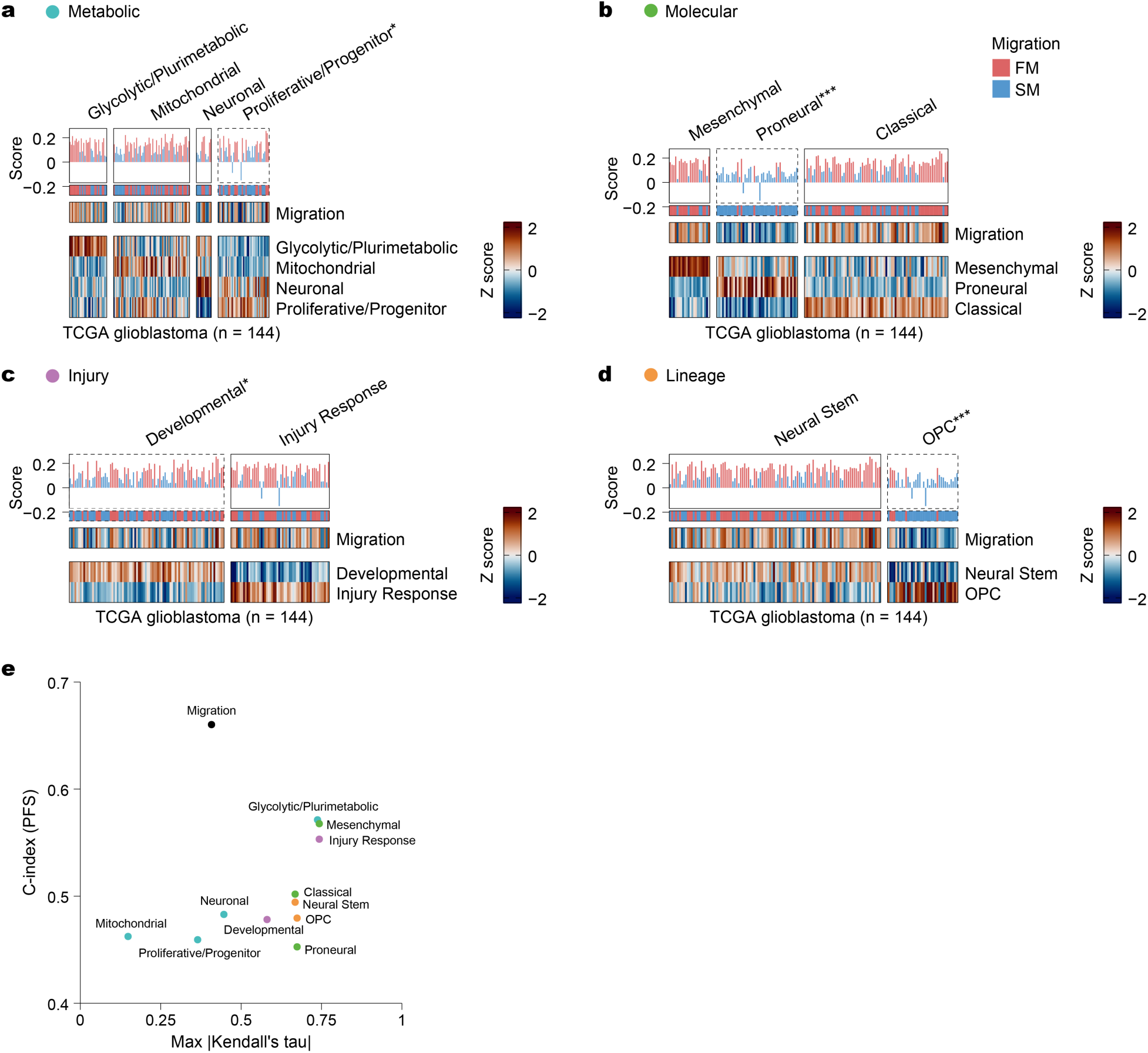
The migration signature retains prognostic value across diverse glioblastoma subtypes. **a–d,** Distributions of migration scores (top) and subtype scores (bottom) for the TCGA glioblastoma RNA-seq cohort (n = 144, IDH1-mutant excluded) clustered by established Metabolic **(a)**, Molecular **(b)**, Injury **(c)**, and Lineage **(d)** classification systems. Dashed boxes and asterisks denote specific subtypes whose median migration score differs significantly from the cohort-wide median (ns ≥ 0.05, * < 0.05, ** < 0.01, and *** < 0.001, two-sided Wilcoxon rank-sum test). **e,** Prognostic performance versus signature uniqueness. Scatter plot comparing the c-index for progression free survival (PFS) against the maximum absolute Kendall’s tau (representing overlap with other signatures) for the migration signature (black) and each subtype in **a–d**. The migration signature isolates to the top-left region, indicating it is both unique (low max tau) and prognostic (high c-index) compared to other signatures.

**Extended Data Fig. 9 (related to Fig. 5):**
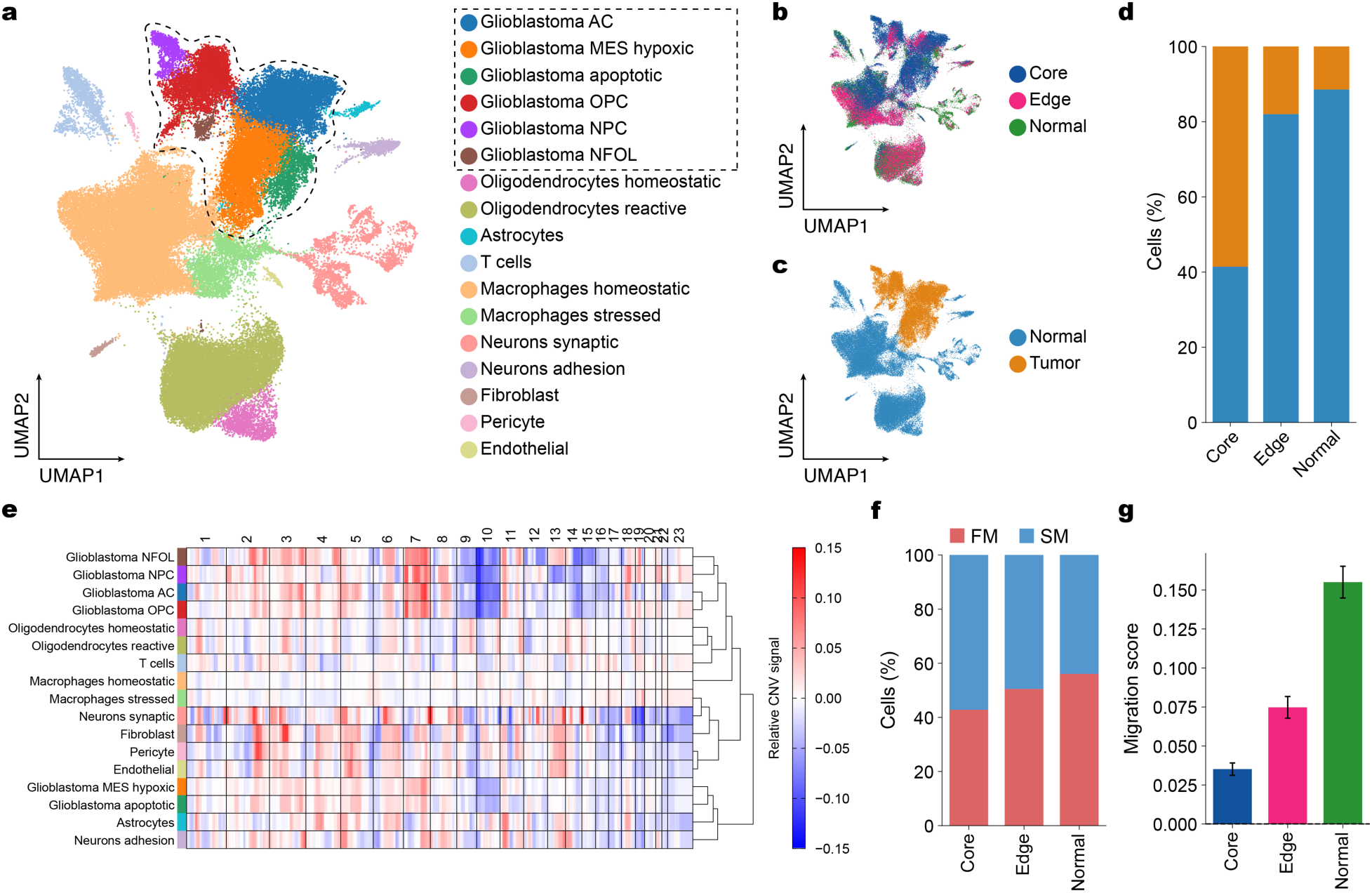
Single-cell transcriptomics map the spatial landscape of glioblastoma. **a,** UMAP embedding of the complete snRNA-seq dataset, annotated by major cell lineages. The dashed line outlines the glioblastoma cell clusters. **b, c,** UMAP embeddings colored by tumor sampling region **(b)** and tumor versus normal classification **(c)**. **d,** Stacked bar plot displaying the proportion of tumor (orange) and non-tumor (blue) cells within each anatomical region, confirming the expected decrease in tumor cells and increase in normal cells from core to edge to normal. **e,** Heatmap of inferred copy number variations (CNVs). Rows represent cell types and columns represent chromosomes. Glioblastoma clusters have the characteristic gain of chromosome 7 and loss of chromosome 10, distinguishing them from non-malignant cells. **f,** Composition of cells classified as Slow-Migrating (SM) (blue) or Fast-Migrating (FM) (red) across tumor regions. **g,** Average migration score by spatial region. Data are mean ± s.e.m.

**Extended Data Fig. 10 (related to Fig. 5):**
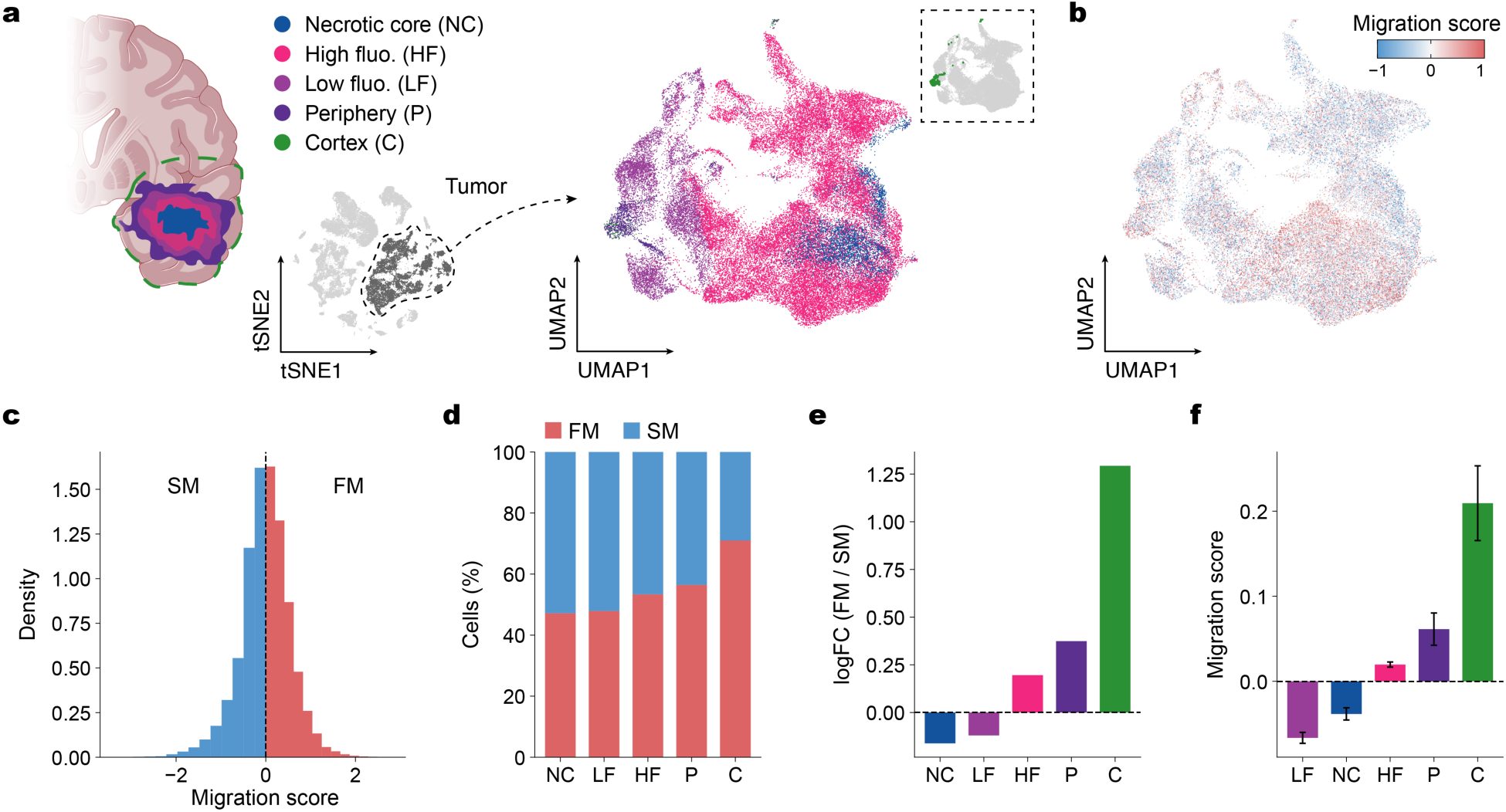
Independent validation of the migration signature in a 5-ALA, region-annotated scRNA-seq dataset confirms enrichment of fast cells at the invasive periphery. **a,** Schematic of the sampling strategy and anatomic labels: cortex (C), periphery (P), low fluorescence (LF), high fluorescence (HF), and necrotic core (NC). Tumor cells were identified (left t-SNE, dashed line outlines tumor cluster) and reanalyzed (right UMAP). Inset (top right) shows the location of cortex cells (green) on the UMAP. **b,** UMAP embedding of tumor cells colored by migration score. Warmer colors indicate higher migration scores (FM-like), cooler colors indicate lower scores (SM-like). **c,** Distribution of migration scores across all tumor cells, showing the threshold (dashed line at 0) used to classify cells as Fast-Migrating (FM) or Slow-Migrating (SM). **d,** Composition of SM (blue) and FM (red) cells within each anatomic site reveals increased fraction of FM cells at the Cortex and Periphery relative to Core. **e,** Log2-fold change (logFC) of the FM/SM cell ratio by anatomic region, highlighting significant enrichment of FM cells in the cortex and periphery. **f,** Average migration score by anatomic site. Data are mean ± s.e.m.

