## Supplemental Figures for "*Ex vivo* glioblastoma migration phenotypes define clinical recurrence and tumor heterogeneity"

**Supplementary Figures**

**Supplementary Figure 1**

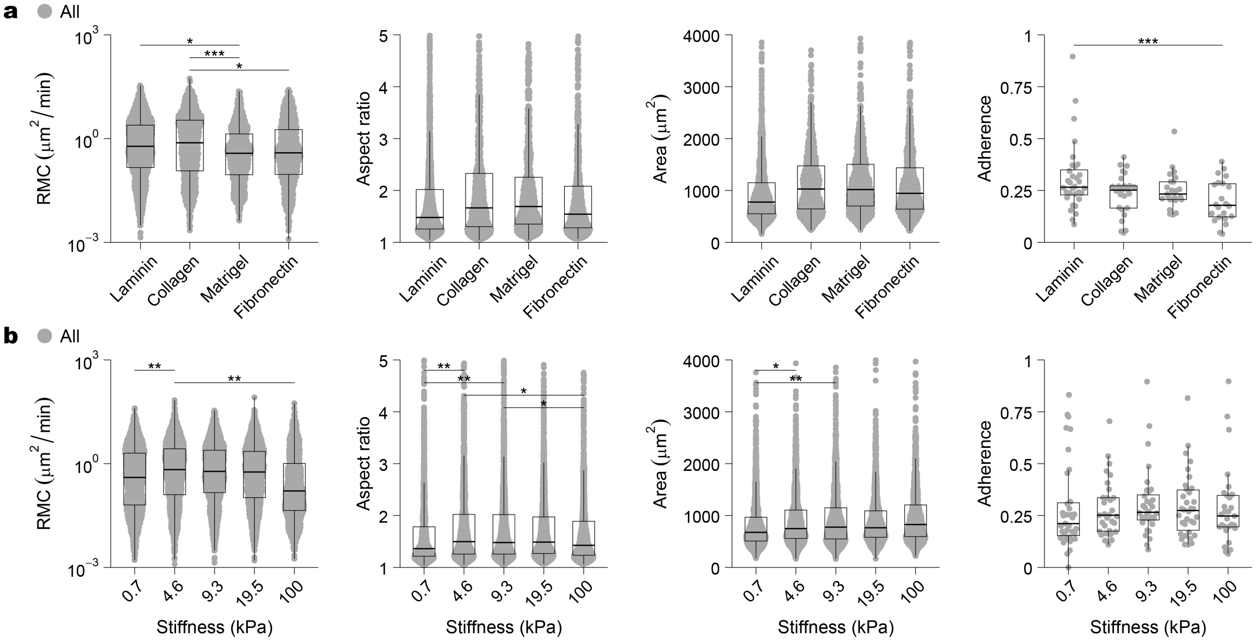


**Supplementary Fig. 1 (related to Fig. 1): Cell-level migration features across substrate ECMs and stiffness levels. a, b,** Distributions of cell-level migration metrics (random motility coefficient (RMC), aspect ratio, area) and patient-level fractional adherence across ECM ligands **(a)** or stiffness levels **(b)**. Data are displayed as box-and-whisker plots overlaid with cell-level data points. Horizontal bars indicate significant pairwise differences determined by linear mixed-effects models accounting for nested random effects (cells within patients) with Tukey-adjusted post-hoc comparisons. Significance: ns ≥ 0.05, * < 0.05, ** < 0.01, and *** < 0.001.

### Supplementary Figure 2

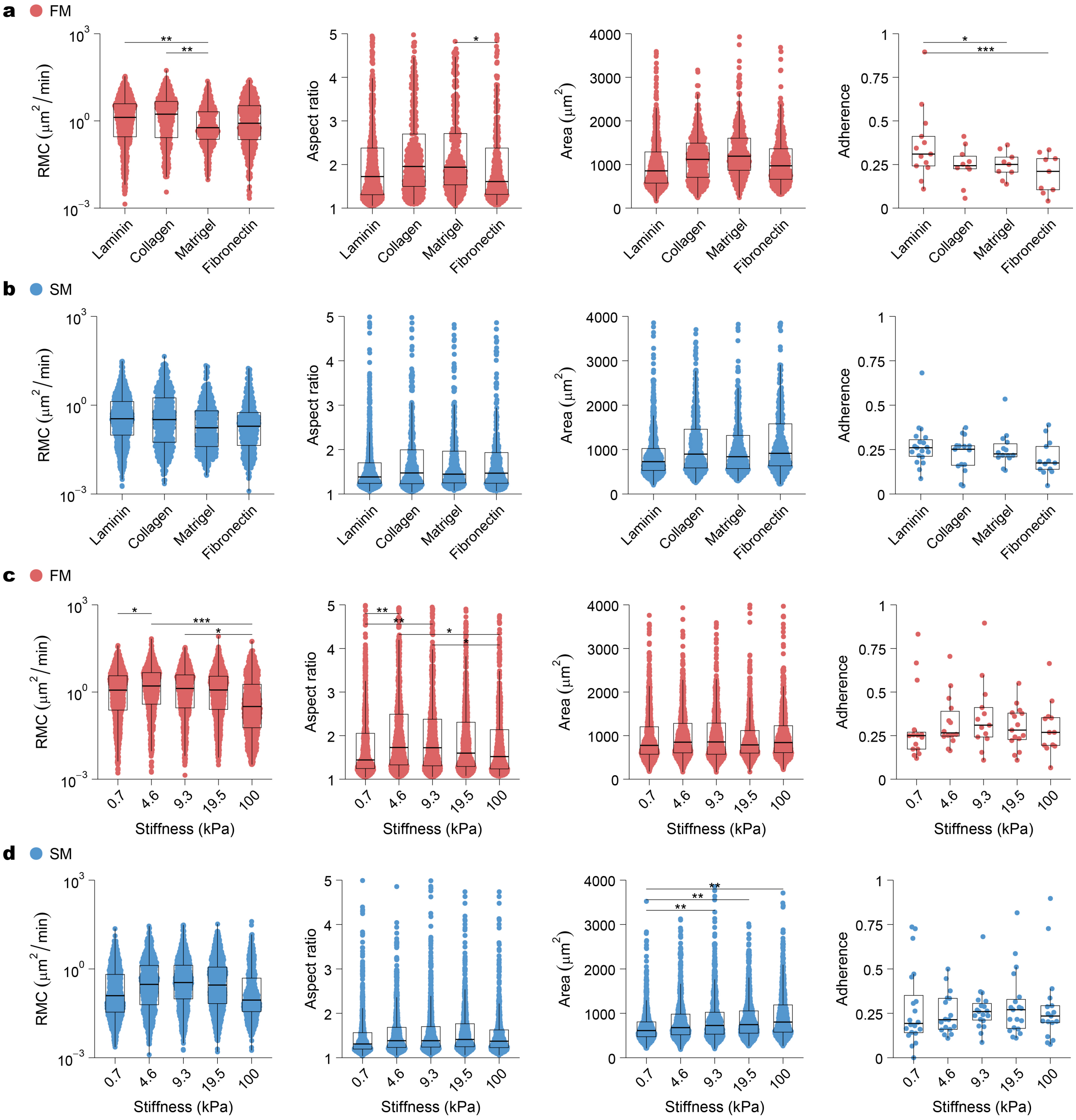


**Supplementary Fig. 2 (related to Fig. 1): Cell-level migration features across substrate ECMs and stiffness levels stratified by FM and SM groups. a–d**, Distributions of cell-level migration metrics (random motility coefficient (RMC), aspect ratio, projected cell area) and patient-level fractional adherence for FM **(a, c)** and SM **(b, d)** groups, analyzed across ECM ligands **(a, b)** or stiffness levels **(c, d)**. Data are displayed as box-and-whisker plots overlaid with cell-level data points. Horizontal bars indicate significant pairwise differences determined by linear mixed-effects models accounting for nested random effects (cells within patients) with Tukey-adjusted post-hoc comparisons. Significance: ns ≥ 0.05, * < 0.05, ** < 0.01, and *** < 0.001.

### Supplementary Figure 3

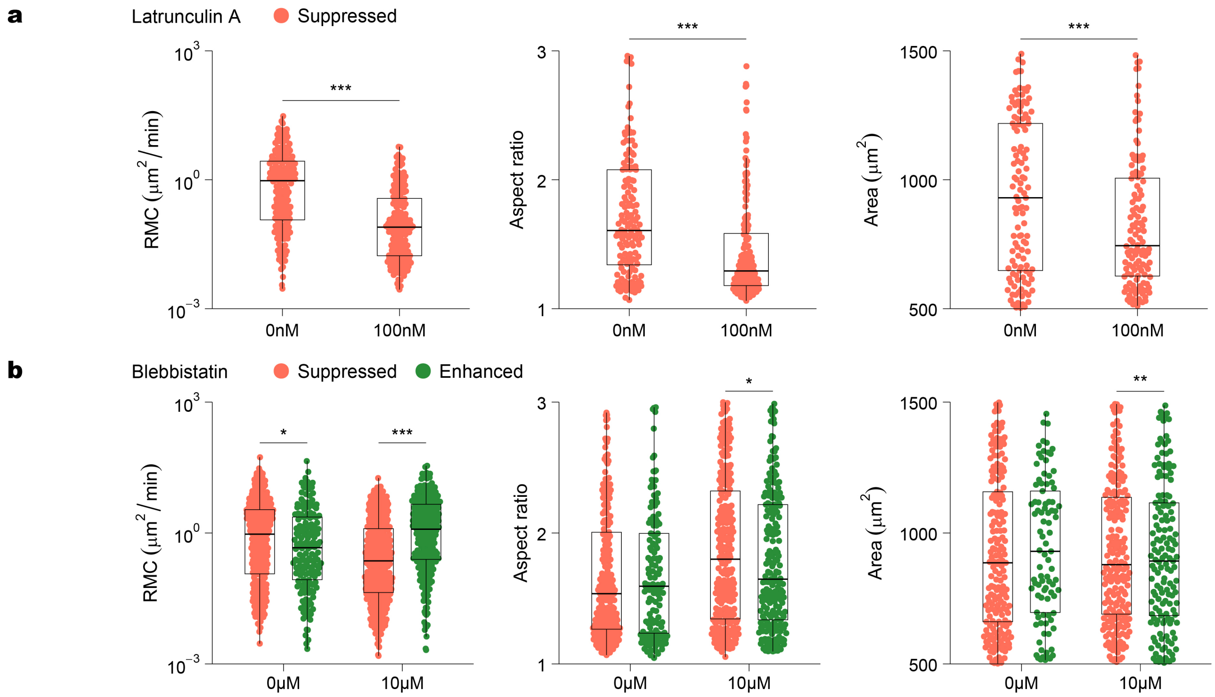


**Supplementary Fig. 3 (related to Fig. 2): Cell-level migration features in response to actin and myosin inhibition. a, b,** Cell-level distributions of random motility coefficient (RMC), aspect ratio, and area following treatment with 100 nM latrunculin A **(a)** or 10 μM blebbistatin **(b)**. Cells in **b** are stratified into blebbistatin-enhanced (green) and blebbistatin-suppressed (orange) groups based on change in RMC (greater or less than zero respectively). Data are displayed as box-and-whisker plots overlaid with cell-level data points. Horizontal bars indicate significant differences determined by two-sided Wilcoxon rank-sum tests. Significance: ns ≥ 0.05, * < 0.05, ** < 0.01, and *** < 0.001.

### Supplementary Figure 4

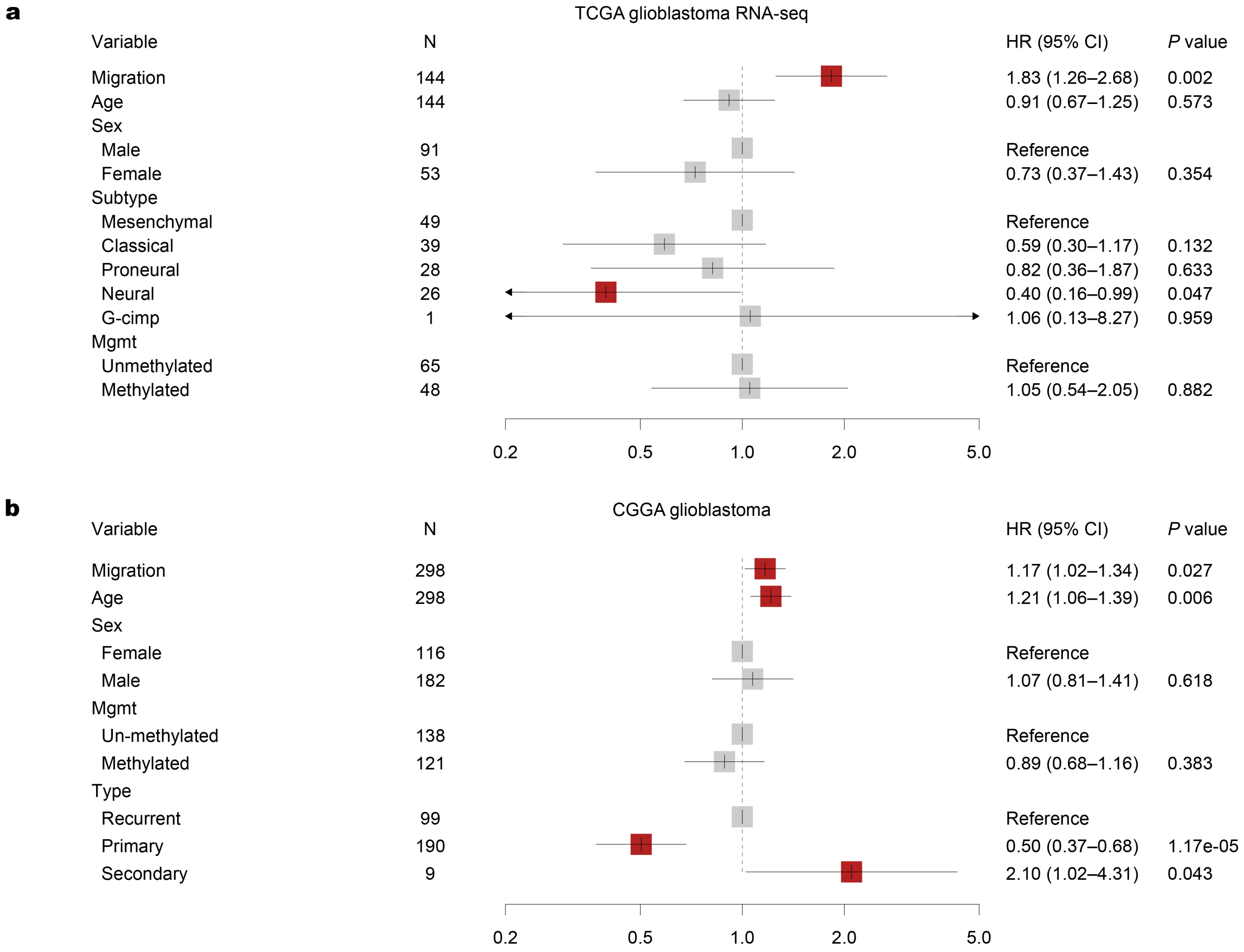


**Supplementary Fig. 4 (related to Fig. 3): Extended prognostic performance of the migration signature. a, b,** Forest plots summarizing hazard ratios (HRs) derived from multivariate (right) Cox proportional-hazards models evaluating PFS in the TCGA glioblastoma RNA-seq cohort **(a)** and OS in the CGGA glioblastoma cohort **(b)**. For TCGA, the multivariate analysis was adjusted for molecular subtype (G-CIMP, Neural, Proneural, and Mesenchymal), MGMT promoter methylation status, sex, and age. For CGGA, the multivariate analysis was adjusted for tumor type (primary, secondary, recurrent), MGMT promoter methylation status, sex, and age. Dots represent HR estimates and horizontal bars indicate 95% confidence intervals. A high migration score (e.g. fast) is associated with significantly increased risk of recurrence and death. In the forest plots, the center of each box represents the estimated hazard ratio, and the whiskers denote the 95% CI. Arrowheads indicate the 95% CI extends beyond the axis limits. CI, Confidence Interval. HR, Hazard Ratio.

### Supplementary Figure 5

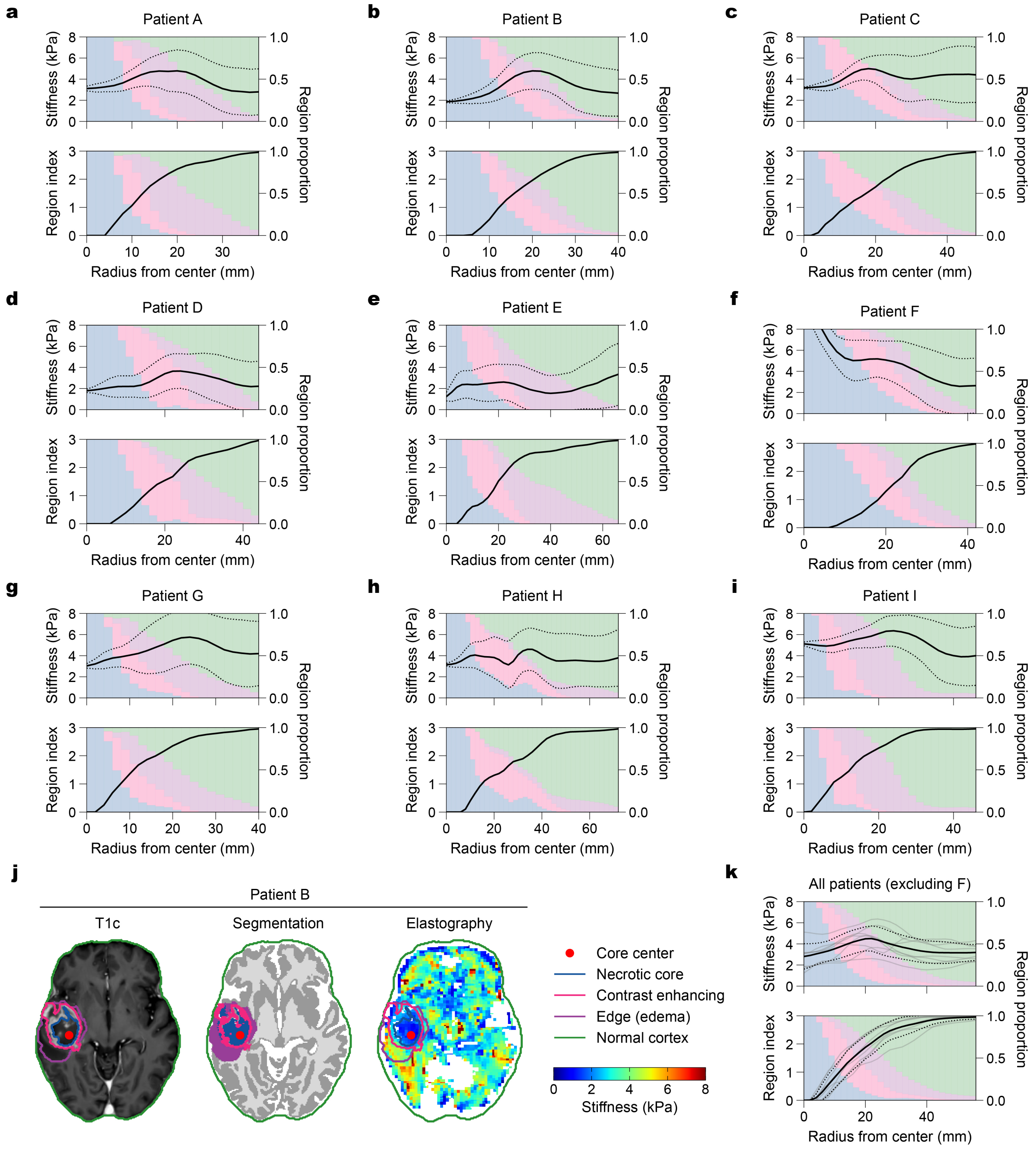


**Supplementary Fig. 5 (related to Fig. 5): Magnetic resonance elastography (MRE) reveals an increase in stiffness at the tumor-brain interface. a–i,** Individual tumor stiffness profiles for nine patients. Upper panels display mean MRE-derived stiffness (solid lines) ± s.d. (dotted lines) calculated using concentric 3D shells expanding from the tumor center. Lower panels show the corresponding mean region index (black line) and regional composition (stacked background colors). Region indices correspond to necrotic core (0), contrast-enhancing tumor (1), edema (2), and normal cortex (3). Background colors indicate the proportion of voxels belonging to each region: necrotic core (gray), contrast-enhancing tumor (pink), edema (blue), and cortex (green). **j,** Representative T1c-weighted MRI, tumor segmentation, and MRE stiffness map from Patient B (corresponding to panel b). **k,** Group-averaged tumor stiffness profile for the cohort (n = 8), excluding Patient F due to an atypical profile (stiff tumor core > 8 kPa). Data are presented as mean ± s.d. The profiles generally demonstrate low stiffness in the tumor core, increasing through the peritumoral transition, peaking at the transition from edema to cortex, and decreasing to normal parenchymal values.

**Supplementary Figure 6**

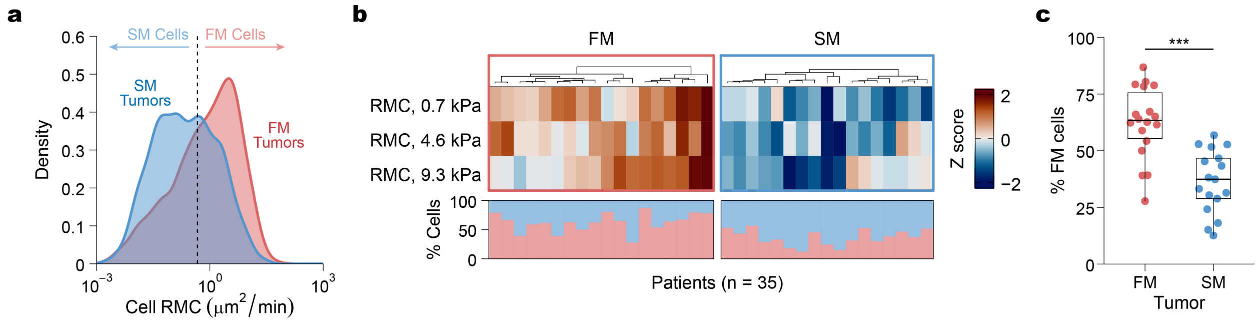


**Supplementary Fig. 6 (related to Fig. 5): Intra-tumoral heterogeneity of single-cell migratory phenotypes in patient-derived glioblastoma. a,** Density distribution of all single-cell RMCs. A threshold defined by the median of all cell RMCs (vertical dashed line) categorizes individual cells into distinct SM (left) or FM (right) subpopulations. Density curves highlight the distribution of cells originating from tumors that were globally classified as Slow-Migration (SM, blue) or Fast-Migrating (FM, red) (as in **Fig. 1b**). **b,** Analysis of intra-tumoral cellular heterogeneity across the patient cohort (n = 35). Top: Heatmap of patient-level hierarchical clustering as in **Fig. 1b**. Bottom: Corresponding stacked bar graphs aligned to each patient, illustrating the intra-tumoral composition as the relative percentage of SM (blue) and FM (red) cells comprising each tumor. **c,** Box plot comparing the percentage of FM cells present within tumors globally classified as SM versus FM. Tumors in the FM patient cohort exhibit a significantly higher intra-tumoral proportion of FM cells ($P$ = 0.00071). Significance: ns ≥ 0.05, * < 0.05, ** < 0.01, and *** < 0.001.

### Supplementary Figure 7

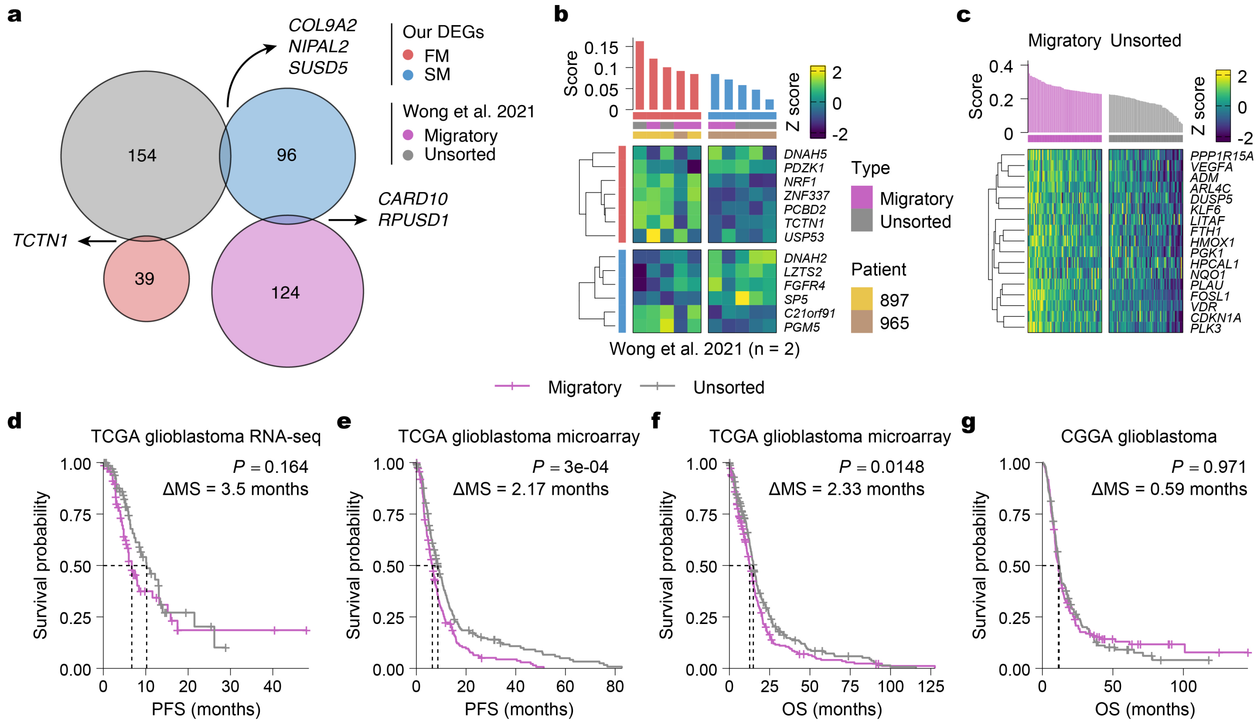


**Supplementary Fig. 7 (related to Fig. 4): The migration signature is distinct from and superior to a previously reported motility signature. a,** Venn diagram illustrating the overlap between DEGs identified in this study and DEGs reported by Wong et al. 2021 [20] (sorted highly motile vs. unsorted cells; 2 patients, triplicate and duplicate samples). Only 3 genes show concordant overlap (*COL9A2*, *NIPAL2*, *SUSD5*). **b,** Heatmap of the 13-gene migration signature from this study applied to RNA-seq data from Wong et al. 2021. Samples (columns) are annotated by type (migratory or unsorted) and patient (897 or 965). Our migration signature mainly stratifies samples by patient instead of type. **c,** Distribution of enrichment scores (top) and expression heatmap (bottom) of the 17 genes defined in the high-motility signature by Wong et al. 2021, applied to the TCGA glioblastoma RNA-seq cohort (n = 144, IDH1-mutant excluded) stratified by median score. Columns represent individual patients and rows represent Z scored expression. **d,** Kaplan-Meier survival analysis of the TCGA glioblastoma RNA-seq cohort stratified by the Wong et al. 2021 signature (as in **c**) shows no significant difference in PFS. **e, f,** Kaplan-Meier analysis of the TCGA glioblastoma microarray cohort (which the Wong et al. 2021 was optimized on) stratified by the Wong et al. 2021 signature shows significantly lower PFS **(e)** and OS **(f)** in the migratory group, as expected. **g,** Kaplan-Meier survival analysis of the CGGA glioblastoma cohort stratified by the Wong et al. 2021 signature shows no significant difference in OS. *P* values were determined by two-sided log-rank test. ΔMS, difference in median survival months.
